# Structural insights into an evolutionary turning-point of photosystem I from prokaryotes to eukaryotes

**DOI:** 10.1101/2022.01.03.474851

**Authors:** Koji Kato, Ryo Nagao, Yoshifumi Ueno, Makio Yokono, Takehiro Suzuki, Tian-Yi Jiang, Naoshi Dohmae, Fusamichi Akita, Seiji Akimoto, Naoyuki Miyazaki, Jian-Ren Shen

## Abstract

Photosystem I (PSI) contributes to light-conversion reactions; however, its oligomerization state is variable among photosynthetic organisms. Herein we present a 3.8-Å resolution cryo-electron microscopic structure of tetrameric PSI isolated from a glaucophyte alga *Cyanophora paradoxa*. The PSI tetramer is organized in a dimer of dimers form with a C2 symmetry. Different from cyanobacterial PSI tetramer, two of the four monomers are rotated around 90°, resulting in a totally different pattern of monomer-monomer interactions. Excitation-energy transfer among chlorophylls differs significantly between *Cyanophora* and cyanobacterial PSI tetramers. These structural and spectroscopic features reveal characteristic interactions and energy transfer in the *Cyanophora* PSI tetramer, thus offering an attractive idea for the changes of PSI from prokaryotes to eukaryotes.

## Introduction

Oxygenic photosynthesis converts light energy into chemical energy and releases molecular oxygen from water, which provides the energy required for sustaining most life activities as well as oxygen needed for all aerobic life on the earth^1^. The light-driven energy conversion reactions are performed by two multi-subunit pigment-protein complexes, photosystem I and photosystem II (PSI and PSII, respectively). Among them, PSII organizes mainly into a dimer throughout photosynthetic organisms from prokaryotes to eukaryotes^2,3^, whereas the structural organization of PSI is significantly diversified among the photosynthetic organisms^4–22^. While prokaryotic cyanobacteria have either trimeric or tetrameric PSI^4–6,9,15–17,20,22^, eukaryotic organisms possess mainly monomeric PSI^7,8,10–14,18,19,21^. Structures of PSI from prokaryotic to eukaryotic organisms have been solved by X-ray crystallography and cryo-electron microscopy (cryo-EM)^4,7–22^, which revealed that the main part of the PSI core is well conserved from prokaryotes to eukaryotes. However, there are differences in the subunit composition and pigment arrangement of the PSI core, reflecting the changes of the PSI core during evolution^23^. Importantly, most of the cyanobacteria do not contain trans-membrane light-harvesting complexes (LHCs), except some specific species of cyanobacteria that contain prochlorophyte chlorophyll (Chl) *a*/*b*-binding proteins^24,25^ and iron-stress-induced-A proteins expressed under iron-deficient conditions^26–31^. Instead, cyanobacteria mainly use water-soluble phycobilisome proteins attached to the stromal side of the membrane as their light-harvesting antennas^32^. On the other hand, eukaryotic PSI core is surrounded by various numbers of trans-membrane LHCS^7,8,10–14,18,19,21^, which makes the PSI core a monomer and prevents it from forming trimers or tetramers. This is one of the major differences found in the PSI structure between prokaryotes and eukaryotes.

*Cyanophora paradoxa* (hereafter referred to as *Cyanophora*) is a glaucophyte alga that is thought to be an ancient eukaryotic alga evolved from prokaryotes, because it has a characteristic chloroplast termed cyanelle^33^. This was corroborated by 16S and 18S rRNA-based phylogenetic analysis showing that the cyanelle is evolutionary very close to cyanobacteria^34,35^. The photosystems of cyanelles use phycobilisomes as their light-harvesting antennas and do not have trans-membrane LHCs. PSI in *Cyanophora* was first found to exist as a monomer^36^, but later PSI tetramers were also found in native membranes by blue-native polyacrylamide electrophoresis (BN-PAGE)^5^. The structure of tetrameric PSI core has been determined from a cyanobacterium *Anabaena* sp. PCC 7120 (hereafter referred to as *Anabaena*)^15–17^, and their excitation-energy-transfer processes have also been observed^15,37^. This raises an interesting question as to whether the *Cyanophora* PSI tetramer has a similar structure and excitation-energy-transfer processes with those of the *Anabaena* PSI tetramer. However, the structural and excitation-energy-transfer properties of the *Cyanophora* PSI tetramer has not been reported yet.

Here we solved a 3.8-Å resolution structure of the PSI tetramer isolated from *Cyanophora* by single-particle cryo-EM analysis. The PSI tetramer showed unique monomer-monomer interactions that are entirely different from the *Anabaena* PSI tetramers^15–17^. Excitation-energy transfer of the *Cyanophora* PSI tetramer is also different from that of the *Anabaena* PSI tetramer. These results illustrate that the *Cyanophora* PSI is in the middle of a shift from oligomers to monomers in this primitive eukaryotic alga during evolution from prokaryotes to eukaryotes.

## Results

### Overall structure of the *Cyanophora* PSI tetramer

The PSI-tetrameric cores were purified from *Cyanophora* as described in the Methods section. Biochemical and spectroscopic analyses showed that this complex is functional and intact (Supplementary Fig. 1). To determine the structure of the PSI tetramer, cryo-EM images of the wild-type PSI tetramer were obtained by a Talos Arctica electron microscope operated at 200 kV. After data processing of the resultant images by RELION (Supplementary Fig. 2, 3, and Supplementary Table 1), a final density map of the wild-type PSI tetramer was obtained with a C2 symmetry at a resolution of 4.0 Å, based on the “gold standard” Fourier shell correlation (FSC) = 0.143 criterion (Supplementary Fig. 2–6 and Supplementary Table 1). However, a dimeric form of the PSI was found in a part of the particles in the process of 3D reconstruction (Supplementary Fig. 2), which suggests that the *Cyanophora* PSI tetramer is labile during either storage of PSI or cryo-grid preparations for cryo-EM analysis.

To suppress the sample dissociation of the PSI tetramer, we employed the GraFix technique^38^ for the preparation of the PSI tetramer, which cross-links protein subunits by glutaraldehyde before freezing of the sample for cryo-EM. The cryo-EM images of the GraFix-treated, cross-linked PSI tetramer (hereafter termed GraFix PSI tetramer) were obtained by the same Talos Arctica electron microscope at 200 kV. After data processing of the resultant images by RELION (Supplementary Fig. 7, 8, and Supplementary Table 1), a final density map of the GraFix PSI tetramer was obtained with a C2 symmetry at a resolution of 3.8 Å (Fig. 1A, and Supplementary Fig. 8, 9). The cryo-EM density map of the GraFix PSI tetramer showed features of well-resolved side chains of most amino acid regions and co-factors.

**Fig. 1.**
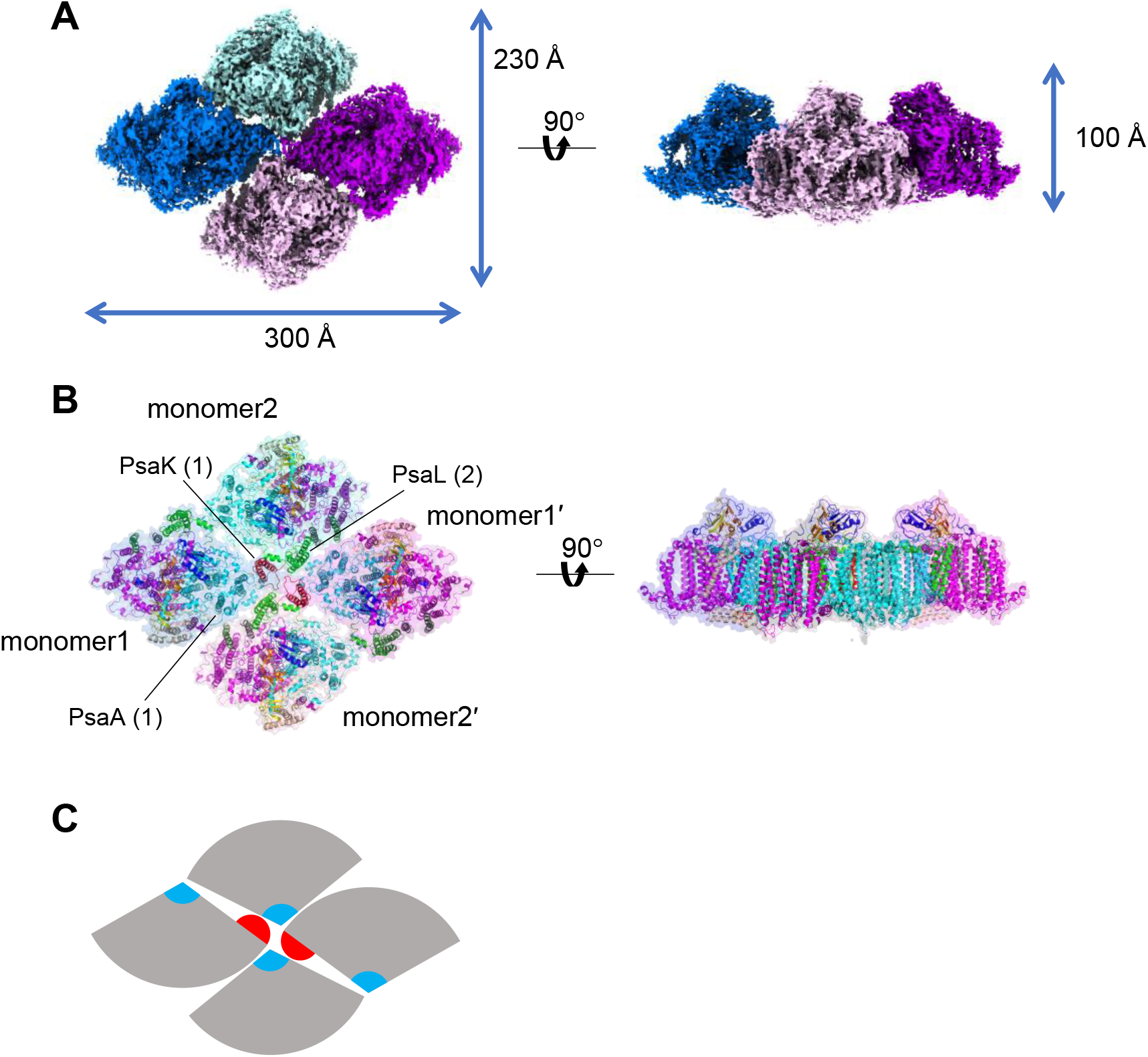
Overall structure of the GraFix PSI tetramer. **A**, The 3D cryo-EM density map of the PSI tetramer viewed along the membrane normal from the stromal side (left) and its side view (right). **B**, The structures of the PSI tetramer viewed along the membrane normal from the stromal side (left) and its side view (right). **C**, Schematic diagram of the PSI tetramer. PsaK and PsaL are shown in red and cyan, respectively.

The four monomers of a PSI tetramer were denoted as monomer1, monomer2, monomer1’, and monomer2’, respectively (Fig. 1B, and Supplementary Fig. 2–6). The dimeric PSI unit is organized by interactions between monomersl(l’) and 2(2’), forming a pseudo-two-fold symmetry. This reflects that the tetramer is assembled by a dimer of dimers, which are designated as monomer1/2-dimer and monomer1’/2’-dimer, respectively (Fig. 1). Examination of the subunits in the tetramer showed that PsaK is present in monomer1 and monomer1’ but absent in both monomer2 and monomer2’ (Fig. 1, 2). Mainly three PSI subunits, PsaA, PsaK, and PsaL, contribute to the interactions between different monomers; in particular, PsaK from one monomer is tightly associated with PsaL from the adjacent monomer at the center of the tetramer (Fig. 1B, 1C). These interactions have not been observed in other structures of PSI trimers and tetramers^4,9,15–17,20,22^. In particular, the *Anabaena* PSI tetramer has four PsaL at the center of the tetramer and four PsaK at the edge of the tetramer^15–17^, and no interactions between PsaL and PsaK are found in the *Anabaena* PSI tetramer (Supplementary Fig. 10). In the *Cyanophora* PSI tetramer, two monomers denoted as monomer1/1’ in Fig. 1 are rotated approximately 90° relative to its counterpart in the *Anabaena* PSI tetramer, resulting in the direct interaction of PsaL of one monomer (monomer2/2’) with PsaK of the adjacent monomer (monomer1/1’) at the center of the *Cyanophora* PSI tetramer. On the other hand, at the edge of the tetramer, PsaK of one monomer (monomer2/2’) is too close to PsaL from the adjacent monomer (monomer1/1’), leading to the loss of PsaK, which occurs in monomer2 and monomer2’ (Fig. 1, and Supplementary Fig. 10). These results demonstrate that the *Cyanophora* PSI tetramer is assembled by novel and unique interactions totally different from the *Anabaena* PSI tetramer.

**Fig. 2.**
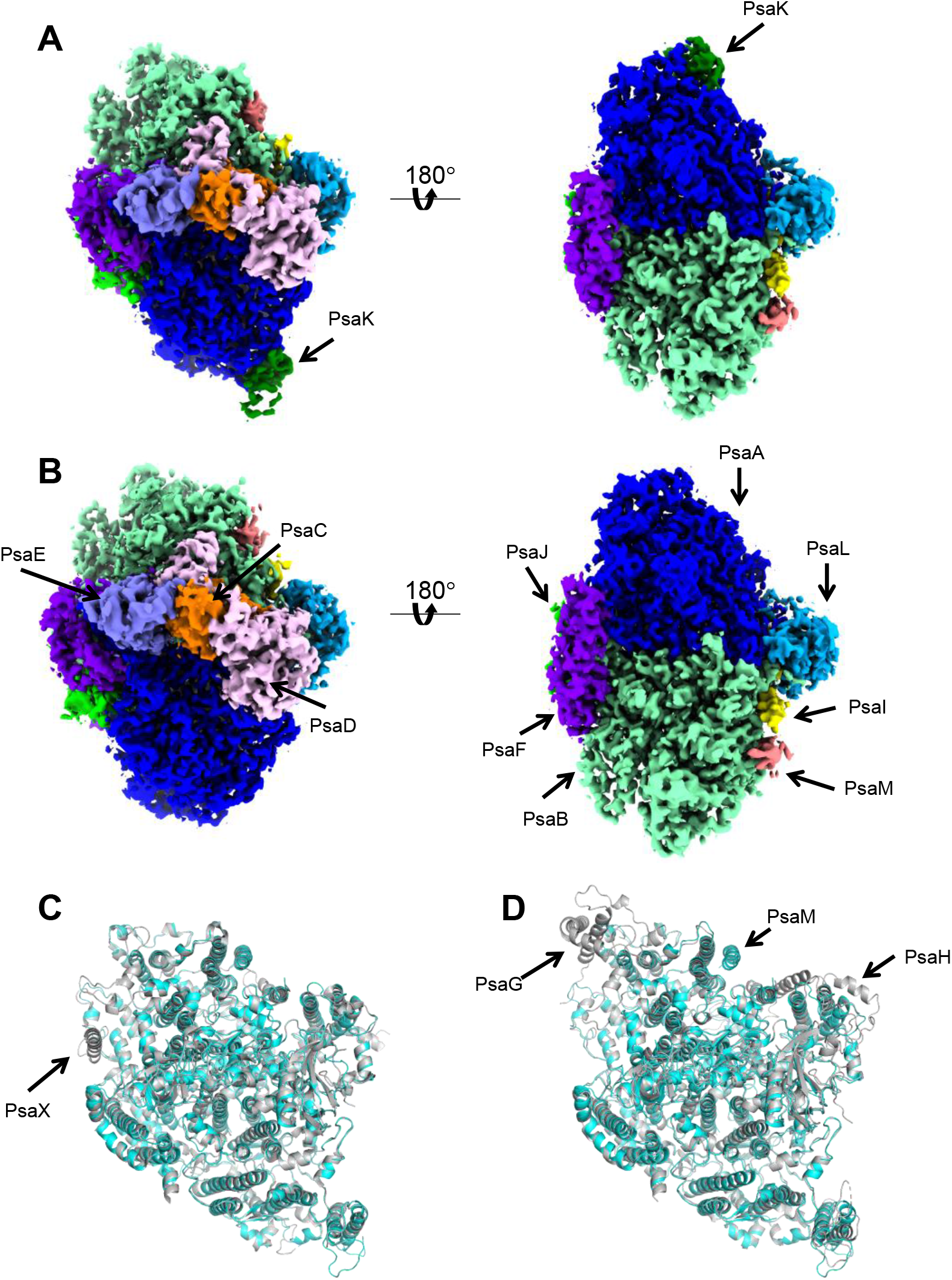
Structure of the wild-type PSI monomer. **A,** The 3D map of monomer1 viewed along the membrane normal from the stromal (left) and lumenal sides (right). **B,** The 3D map of monomer2 viewed along the membrane normal from the stromal (left) and lumenal sides (right). **C,** Superposition of the *Cyanophora* PSI monomer1 (cyan) with a PSI-monomer unit from *T. elongatus* (gray), viewed along the membrane normal from the stromal side. **D,** Superposition of the *Cyanophora* PSI monomer1 (cyan) with a PSI-monomer unit from *P. sativum* (gray), viewed along the membrane normal from the stromal side. The subunits specific to each organism are labeled.

### Structure of the PSI monomers

For the accurate model building, we performed focused 3D classifications using masks covering each monomeric unit (monomer1 and monomer2). The final cryo-EM density maps of wild-type PSI monomer1 and monomer2 were obtained with a C1 symmetry at resolutions of 3.3 Å and 3.2 Å, respectively (Fig. 2, Supplementary Fig. 2, 3, and Supplementary Table 1). Monomer1 contains well-known 8 membrane-spanning subunits (PsaA, PsaB, PsaF, PsaI, PsaJ, PsaK, PsaL, and PsaM) and three stromal subunits (PsaC, PsaD, and PsaE) (Fig. 2). The subunit composition of monomer2 is comparable to that of monomer1 except that PsaK is lacking. The structure of a PSI-monomer unit within the tetramer is similar to that in the cyanobacterial and plant PSI cores (Fig. 2C, 2D), except that the cyanobacterial PSI contains an additional subunit PsaX, whereas the higher plant PSI has additional PsaG and PsaH subunits but without PsaM (Fig. 2C, 2D). The *psaG*, *psaH*, and *psaX* genes are not found in the genome of *Cyanophora*^39^, indicating the loss of PsaX and prior to the acquisition of PsaG and PsaH in this primitive eukaryote. There are two copies of the *psaA* and *psaB* genes in *Cyanophora*, namely, *psaA1*/*psaA2* and *psaB1*/*psaB2*; however, all of the PsaA and PsaB subunits in the tetramer structure is identified as the gene products of *psaA1* and *psaB1*. The other nine subunits have only one gene.

The cofactors identified in the monomer1 and monomer2 of the tetramers are summarized in Supplementary Table 2. Monomer1 has 84 Chls *a*, 19 *β*-carotenes, 3 [4Fe-4S] clusters, 2 phylloquinones, and 3 lipid molecules, whereas monomer2 possesses 81 Chls *a*, 19 *β*-carotenes, 3 [4Fe-4S] clusters, 2 phylloquinones, and 3 lipid molecules. The locations of these molecules are similar to those in the prokaryotic and eukaryotic PSI structures^4,7–22^, although the number of Chls differs significantly. Some of the Chls are lost from PsaB, which is likely due to the disordered structure around PsaB and/or dissociation of the cofactors during preparation of the *Cyanophora* PSI cores.

### Interactions between monomer1 and monomer2 within a dimer

Because of the rotation of two monomers in the tetramer (Supplementary Fig. 10), the interactions among monomers of the *Cyanophora* PSI tetramer are clearly different from those of the *Anabaena* PSI tetramer. One of the interfaces, the interface between monomer1 and monomer2 of the *Cyanophora* PSI tetramer, is formed between PsaA/K of monomer1 and PsaA/L of monomer2 at the center of the tetramer, as well as PsaL of monomer1 and PsaA of monomer2 at the edge of the tetramer (Fig. 3A). At the lumenal side, Asn497 of monomer1-PsaA is tightly coupled with Asn497 of monomer2-PsaA at distances of 2.5–2.6 Å (Fig. 3B). Gln88 and Ser89 of monomer1-PsaK interact with Val64, Glu65, Arg70, and Asn71 of monomer2-PsaL at distances of 2.8–3.2 Å (Fig. 3C), at the center of the tetramer. Monomer1-PsaK is also associated with monomer2-PsaL through hydrophobic interactions between Ala109/Pro113 of monomer1-PsaK and Ala27/Val28 of monomer2-PsaL at the stromal side of the center (Fig. 3D), and Ala27/Gly30/Leu31 of monomer1-PsaL interacts with Gly317/Ile318 of monomer2-PsaA through hydrophobic interactions at the stromal side of the edge (Fig. 3E). In addition, many protein-pigment interactions are found in the monomer1 and monomer2 interface.

**Fig. 3.**
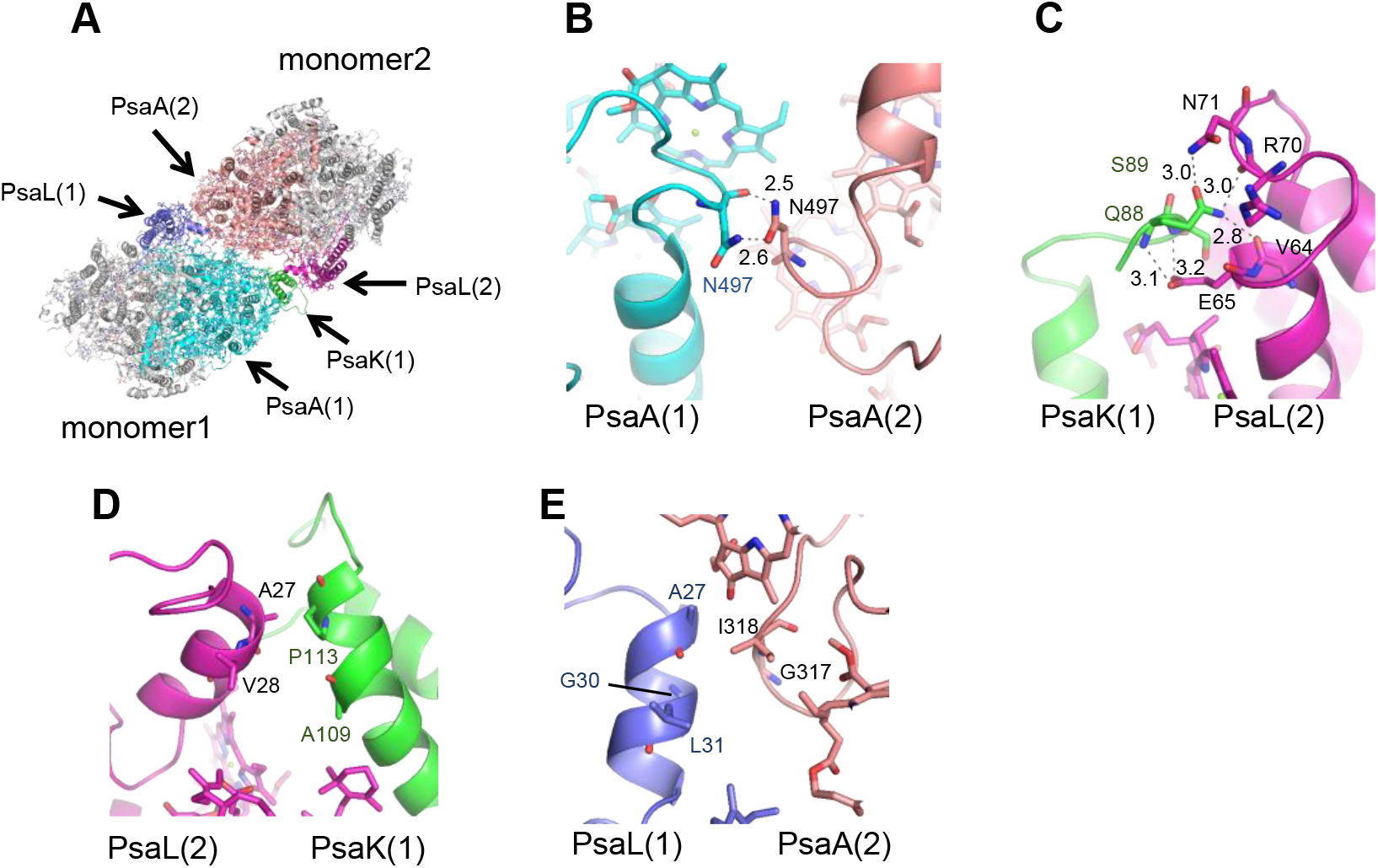
Intermonomer interactions between monomer1 and monomer2. **A,** Structure of a monomer1/2 dimer unit from the PSI tetramer viewed along the membrane normal from the stromal side. **B,** Interactions between monomer1-PsaA and monomer2-PsaA. **C, D,** Interactions between monomer1-PsaK and monomer2-PsaL. **E,** Interactions between monomer1-PsaL and monomer2-PsaA.

### Interactions between monomer1 and adjacent monomer2’

The other interface in the *Cyanophora* PSI tetramer is the interface between monomer1 and adjacent monomer2’, which is formed between PsaA of monomer1 and PsaI/L/M of monomer2’ (Fig. 4A). At the lumenal side, Chl817 of monomer1-PsaA interacts with Asn4 of monomer2’-PsaI at a distance of 2.8 Å (Fig. 4B), Chl816 of monomer1-PsaA is hydrogen-bonded to Tyr139 of monomer2’-PsaL at a distance of 3.1 Å (Fig. 4C), and Tyr160/Ile164 of monomer1-PsaA interact with Phe8/BCR101 of monomer2’-PsaM through hydrophobic interactions (Fig. 4D). On the other hand, no apparent interactions are found between monomer1 and monomer2’ at the stromal side.

**Fig. 4.**
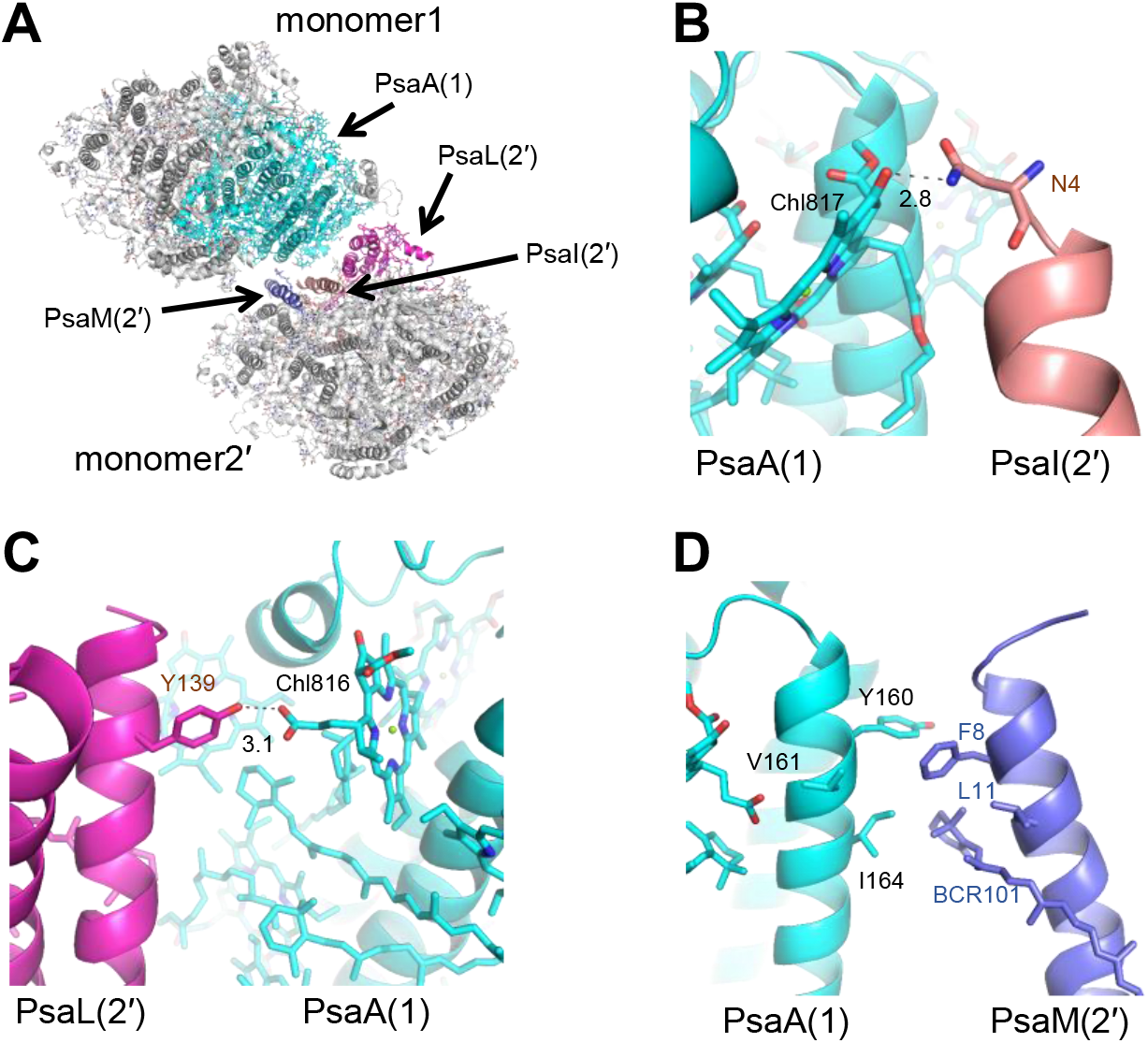
Intermonomer interactions between monomer1 and monomer2’. **A,** Structure of a monomer 1/2’ dimer unit from the PSI tetramer viewed along the membrane normal from the stromal side. **B**, Interactions between monomer1-PsaA and monomer2’-PsaI. **C,** Interactions between monomer1-PsaA and monomer2’-PsaL. **D,** Interactions between monomer1-PsaA and monomer2’-PsaM.

### Interactions at the center of the tetramer

As mentioned above, each monomer is associated at the center of the tetramer through interactions between PsaK of one monomer and PsaL of the adjacent monomer (Fig. 5A). At the stromal side, Asn127 of monomer1(1’)-PsaK interacts with Gln129 of monomer1’(1)-PsaK at a distance of 2.8 Å (Fig. 5B). There are several complicated interactions among monomer1-PsaK, monomer2-PsaL, monomer1’-PsaK, and monomer2’-PsaL (Fig. 5C). The loop structure between Pro125 and Pro131 of monomer1-PsaK interacts with its counterpart from monomer1’-PsaK through hydrophobic interactions. Chl201 in monomer2-PsaL interacts with Pro131 of monomer1-PsaK and Phe126 of monomer1’-PsaK, and Chl201 in monomer2’-PsaL interacts with Pro131 of monomer1’-PsaK and Phe126 of monomer1-PsaK, through hydrophobic interactions.

**Fig. 5.**
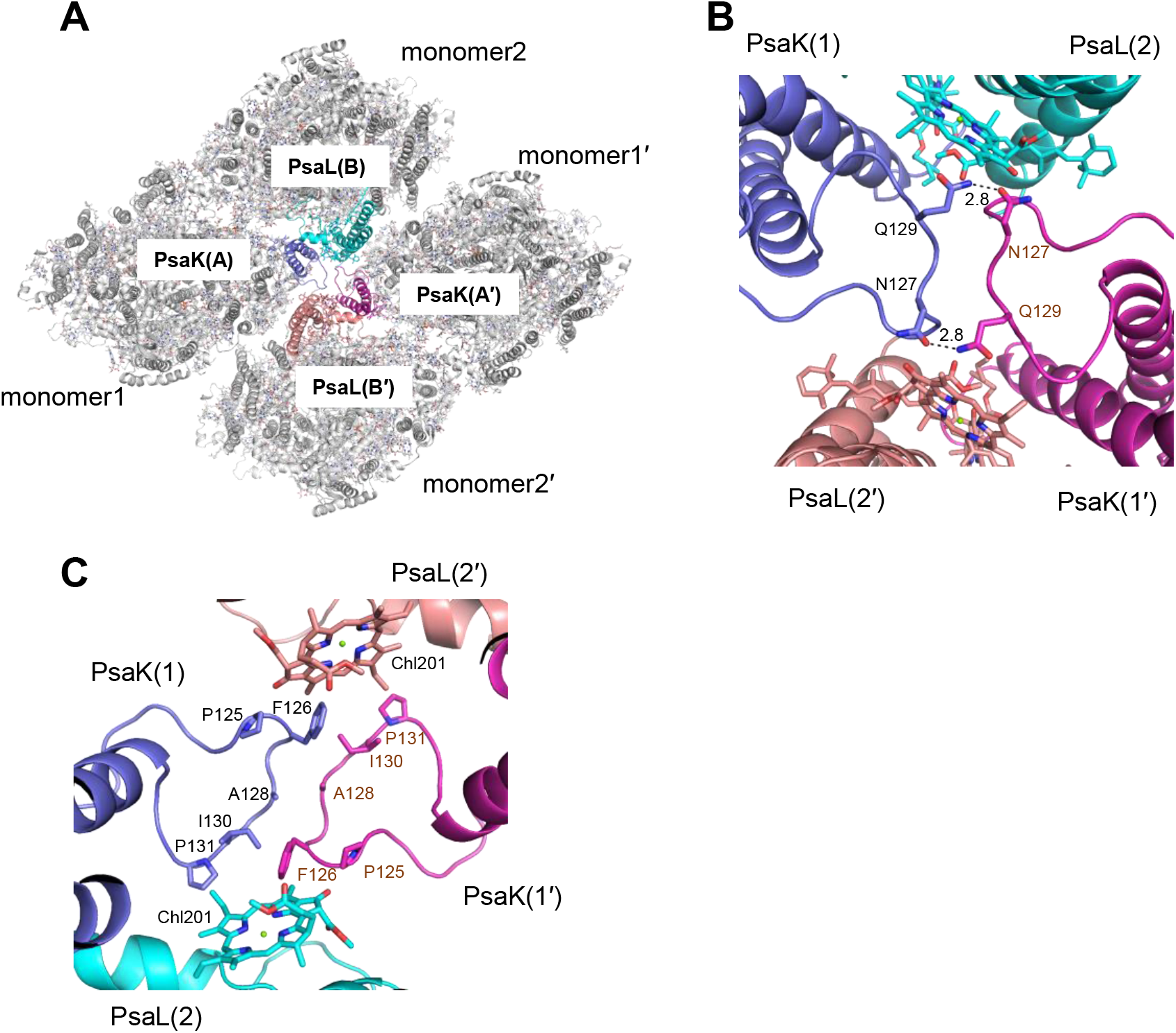
Interactions at the center of the PSI tetramer. **A,** Structures of the PSI tetramer viewed along the membrane normal from the stromal side. **B, C,** Interactions between monomer1-PsaK, monomer2-PsaL, and monomer1’-PsaK, monomer2’-PsaL.

### Pigment-pigment interactions in the tetramer

The location of pigment molecules in monomer1 is similar to that in monomer2, except that three Chls *a* (PsaA-Chl845, PsaB-Chl835, and PsaK-Chl201) in monomer1 are absent in monomer2 (Supplementary Fig. 11, and Supplementary Table 3). The lack of PsaB-Chl835 in monomer2 is likely due to disordered structure around PsaB or to dissociation of the cofactors during preparation of the PSI tetramer, whereas the lack of PsaA-Chl845 and PsaK-Chl201 may be due to the loss of PsaK in monomer2, because they are located at positions near PsaK in monomer1.

We have recently identified characteristic Chl clusters, namely Low1 and Low2, based on structural comparison of PSI trimers among three types of cyanobacteria, *Gloeobacter violaceus* PCC 7421, *Synechocystis* sp. PCC 6803, and *Thermosynechococcus vulcanus* NIES-2134^22^. Low1 is composed of a dimeric Chls, whereas Low2 is composed of a triple Chls^22^. In the *Cyanophora* PSI tetramer, a Chl1A and Chl2A dimer in monomer1 corresponds to Low1 (Supplementary Fig. 12A). On the other hand, monomer1 does not have Low2 because of the absence of Chl1B that is necessary for the formation of Low2 as seen in the PSI structure of *T. vulcanus* (Supplementary Fig. 12B). Monomer2 also has Low1 but not Low2 (Supplementary Fig. 12C, 12D). Different from the Low2 site in monomer1, monomer2 does not have Chl2B as well as Chl1B. We have suggested that Low1 and Low2 are related to fluorescence peaks at around 723 and 730 nm, respectively, in the fluorescence-emission spectra of PSI trimers from the three types of cyanobacteria^22^. In the *Cyanophora* PSI tetramer, the fluorescence-emission spectrum showed a peak at around 722 nm but lacked the 730 nm peak (Supplementary Fig. 1D), which supports that the *Cyanophora* PSI tetramer has Low1 but not Low2. However, we have to point out that the cryo-EM map around the Low2 site in monomer2 has a rather lower quality.

There are numerous pigment-pigment interactions among the monomers within the PSI tetramer (Supplementary Fig. 13A). Unique association of pigments within the tetramer are found in the interfaces between monomer1(1’) and monomer2(2’) but not between monomer1(1’) and monomer2’(2), at the stromal side. A triply stacked Chl cluster exists in both monomer1/2 and monomer1’/2’ in their interface, which are composed of Chl823/824/846 (Supplementary Fig. 13B). The edge-to-edge distances among the Chl clusters are in the range of 3.8–4.3 Å. These Chl clusters thus may have lower energy levels. Two *β*-carotenes (BCR849 and BCR852) are close to these Chl clusters at distances of 3.6–4.6 Å (Supplementary Fig. 13B), suggesting a close interaction among these Chls and *β*-carotenes, which may enable either highly efficient energy transfer or quenching between Chls and *β*-carotenes. In addition, PsaL-BCR205 at the stromal side and PsaM-BCR101 at the lumenal side of monomer2’(2) have their chains inserted into the region of monomer1(1’), which may mediate either energy transfer or quenching between PsaM-BCR101/PsaL-BCR205 in monomer2’(2) and PsaA-Ch1808/Ch1845 of monomer1(1’), respectively (Supplementary Fig. 13A, 13C, 13D). On the other hand, Chls in different monomers are located very far at the lumenal side, suggesting less energy transfer among the monomers at the lumenal side.

### Excitation-energy-transfer processes of the tetramer

Time-resolved fluorescence (TRF) spectra of the PSI tetramer were measured at 77 K (Fig. 6). The TRF spectra exhibit two fluorescence bands at around 694 and 717 nm just after excitation (0–4.9 ps). Until 130 ps, the 694-nm band is lost, whereas the 717-nm band is slightly shifted to about 719 nm. This suggests either excitation-energy transfer from Chls fluorescing at 694 and 717 nm to those at 719 nm or energy trapping to the reaction-center Chls in PSI. The 719-nm band is gradually shifted to longer wavelengths to 728 nm until 4.6 ns, suggesting energy transfer to low-energy Chls fluorescing at 728 nm. In addition, fluorescence at around 677 nm is observed in the time range of 0.9–4.6 ns, suggesting fluorescence from uncoupled Chls in the PSI-tetramer fraction. These characteristic peaks in the TRF spectra are verified in the steady-state fluorescence spectrum of the tetramer (Supplementary Fig. 1D).

**Fig. 6.**
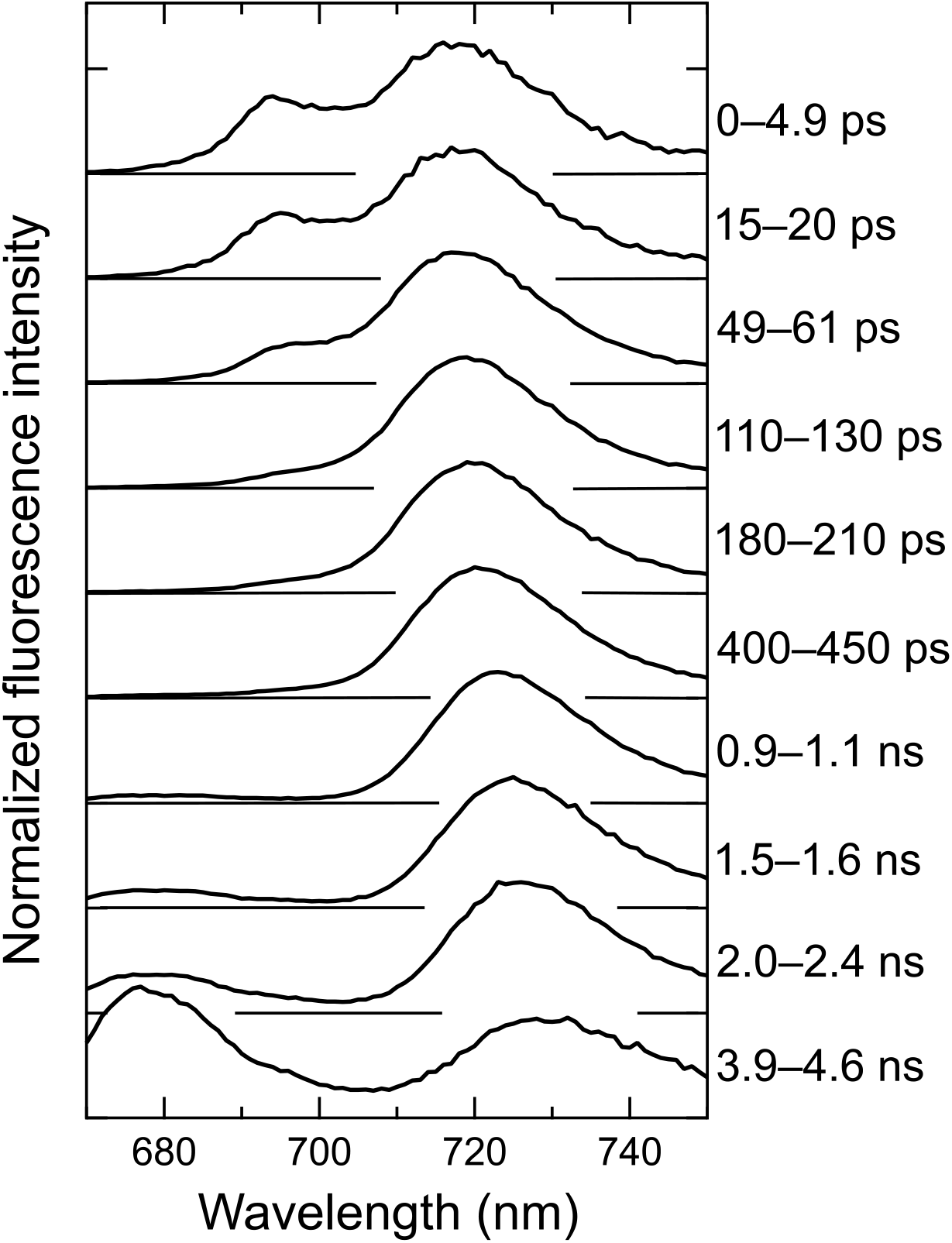
TRF spectra of the *Cyanophora* PSI tetramer. The spectra were measured at 77 K with excitation at 445 nm, and were normalized by the maximum intensity of each spectrum.

## Discussion

### Structural basis for the oligomerization of PSI

The present study showed a completely different arrangement of PSI tetramer of *Cyanophora* compared with that of *Anabaena* PSI tetramer, thereby providing a potential clue as to how the PSI oligomerization state is determined. We compare the structures of the *Cyanophora* PSI tetramer with those of PSI trimer and tetramer in cyanobacteria (Supplementary Fig. 14A). Here we focus on the PsaL subunit because of its significant contribution to the assembly of cyanobacterial PSI^4,9,15–17,20,22,40,41^. Superposition of the *Cyanophora* PSI monomers with cyanobacterial PSI trimer from *Thermosynechococcus elongatus* (PDB: 1JB0, hereafter referred to as *T. elongatus*) showed a steric hindrance by Arg45 and Ile129 of PsaL (Supplementary Fig. 14B, 15). Ile129 is found in *Anabaena* PsaL, and a similar Val129 is found in other cyanobacteria that form PSI trimer (Supplementary Fig. 15). However, an uncharged residue is found to replace Arg45 in cyanobacteria that form either PSI trimer or tetramer (Supplementary Fig. 15). Furthermore, the C-terminus of *Cyanophora* PsaL is shorter than cyanobacterial PsaL (Supplementary Fig. 14C, 15). These factors may make the *Cyanophora* PsaL unable to form interactions required for the trimer formation. In contrast, superposition of a *Cyanophora* PSI monomer with the *Anabaena* PSI tetramer showed no steric hindrance around PsaL as observed above. Interestingly, the N and C-termini of *Cyanophora* PsaL are shorter than those of *Anabaena* PsaL (Supplementary Fig. 14D–14F), both of which are important for the formation of the *Anabaena* PSI tetramer. This suggests that PsaL of *Cyanophora* is unable to support the formation of a tetramer similarly as that observed in *Anabaena*.

We next examined the structure and sequence of PsaK that may differentiate *Cyanophora* from cyanobacteria and eukaryotes. An insertion between 121–131 is found in the *Cyanophora* PsaK sequence which is absent in cyanobacteria (Supplementary Fig. 16). As described above (Fig. 5B, 5C), residues in this region are required for the interaction of PsaK with PsaL and PsaK from other monomers in the *Cyanophora* PSI tetramer. In addition, there are also some changes in the residues of Gln88, Ser89, Ala109, and Pro113 that interact with PsaL from the adjacent monomer (Fig. 3C, 3D). Among these residues, Gln88 and Ser89 are unique to the *Cyanophora* PsaK, whereas Ala109 and Pro113 are changed to similar or different residues, or are conserved, in cyanobacteria (Supplementary Fig. 16). Thus, the insertion of 121–131 and changes of Gln88 and Ser89 in the sequence of *Cyanophora* PsaK may enable the formation of atypical assembly of the PSI tetramer in *Cyanophora* and may cause rotation of two of the four monomers in the *Cyanophora* PSI tetramer relative to the *Anabaena* PSI tetramer. The insertion around residues 121–131 is also found in green algae and higher plants that form PSI monomer. However, the sequence of this region is highly variable, and the eukaryotic PSI has the trans-membrane LHCs to surround the monomeric PSI core. Phylogenetic analyses of typical PSI subunits, PsaA, PsaB, PsaK, and PsaL, showed that while the *Cyanophora* PsaA, PsaB, and PsaL are either grouped within cyanobacteria or between cyanobacteria and eukaryotes, the sequences of *Cyanophora* PsaK do not group with any of the other organisms and form a unique clade consisting of *Cyanophora* only (Supplementary Fig. 17–20). This is good evidence for the uniqueness of PsaK and its role in tetramer formation in *Cyanophora*.

Despite the presence of a large hole in the center of the *Anabaena* PSI tetramer but almost no free space in the center of the *Cyanophora* PSI tetramer, the interactions of each protomer within the *Cyanophora* PSI tetramer are weaker than those within the *Anabaena* PSI tetramer. This is manifested by the fact that the GraFix method has to be used to isolate the stable *Cyanophora* PSI tetramers, in order to prevent dissociation of the tetramer into monomers. The weak interactions among the PSI-monomer units in *Cyanophora* may be caused by large modifications in the sequences of PsaL and PsaK, and may be helpful for dissociation of the PSI cores from oligomers to monomers. Upon complete attachment of PsaK, the tetramers will be broken and transferred to monomers. This leads us to propose a model for the oligomerization of PSI during evolution from cyanobacteria to various eukaryotes, including higher plants. Changes in the sequences of PsaL result in the cyanobacterial-type PSI tetramer, whereas changes in the PsaL and PsaK sequences result in the *Cyanophora-type* PSI tetramer seen only in the eukaryote *Cyanophora*. Furthermore, PsaH is acquired in some eukaryotes, and this subunit, together with the trans-membrane LHC subunits, may inhibit the oligomerization of PSI. Since PsaH and trans-membrane LHCs are not encoded in the *Cyanophora* genome^39^, it is implied that both changes in the sequences of PsaL and PsaK and the lack of PsaH and LHCs contribute to the formation of the unusual tetrameric PSI structure in *Cyanophora*.

The above structural implications allow us to draw a model implying that *Cyanophora* is an intermediate between oxygenic photosynthetic prokaryotes and eukaryotes in the evolutionary processes of oxyphototrophs (Fig. 7). The cyanobacterial PSI cores are mainly trimers or tetramers. The changes in PsaL, especially its N-terminal region and hydrophobic amino acids, together with PsaK, induce atypical interactions responsible for the tetramerization of the *Cyanophora* PSI cores (Supplementary Fig. 14–20). For the evolution of red and green lineages, the *Cyanophora* PSI tetramers are once dissociated into monomers, and then other PSI subunits and membrane-embedded LHCs are bound to the PSI monomers. The additional bindings of PsaG, PsaH, and LHCs inhibit PSI oligomerization in the eukaryotes (Fig. 7). These observations are consistent with the evolutionary point of view that *Cyanophora* is evolved from cyanobacteria having PSI tetramers and subsequently serves as an ancestor for other eukaryotic algae where PSI becomes a monomer.

**Fig. 7.**
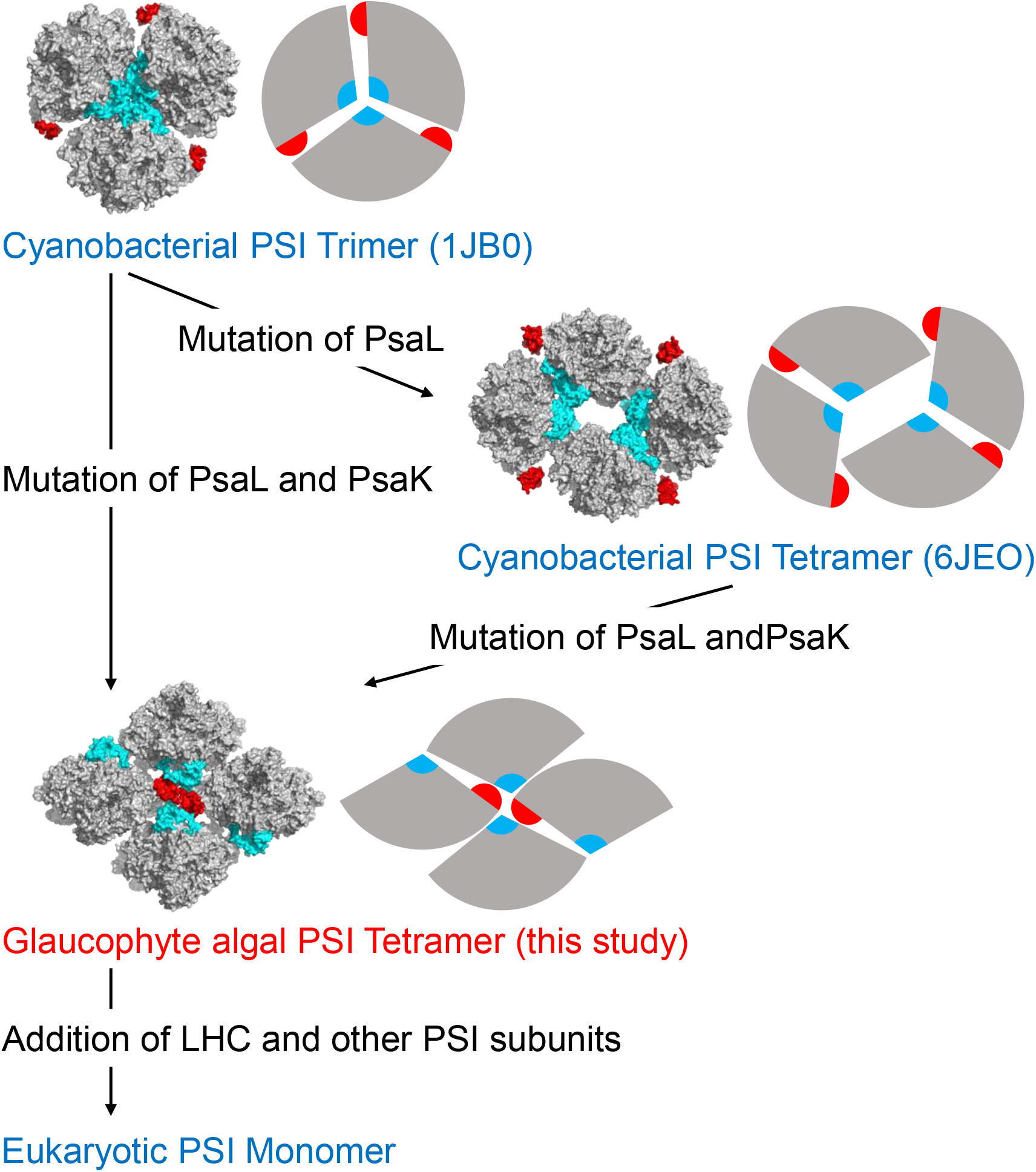
A model for the molecular evolution of PSI assembly. The structures of the PSI oligomer are shown in both surface model and schematic diagram. PsaK and PsaL are shown in red and cyan, respectively.

### Different excitation-energy dynamics of PSI tetramers between *Cyanophora* and *Anabaena*

We have shown excitation-energy-transfer processes of the *Anabaena* PSI tetramer^15,37^. Different from the TRF spectra of the *Cyanophora* PSI tetramer (Fig. 6), the *Anabaena* PSI tetramer showed two fluorescence bands at around 696 and 730 nm with a shoulder at around 715 nm just after excitation (0–4.9 ps)^15^. Among the three fluorescence bands, the fluorescence at around 715 nm decreased until 130 ps in the *Anabaena* PSI tetramer but not in the PSI dimer and monomer^15^, thereby contributing to energy transfer among PSI-monomer units by the Chls fluorescing at 715 nm in the *Anabaena* PSI tetramer. In the *Cyanophora* PSI tetramer, however, the 717-nm fluorescence band still remains at 130 ps (Fig. 6); its relative decrease is not small compared with the decrease in fluorescence at around 715 nm in the TRF spectra of *Anabaena* PSI tetramer. Instead of the 717-nm fluorescence band, the 694-nm band prominently decreases until 130 ps in the *Cyanophora* PSI tetramer (Fig. 6), suggesting significant contribution of energy transfer among PSI-monomer units. These results suggest the clear differences in excitation-energy transfer among PSI-monomer units between *Cyanophora* and *Anabaena* PSI tetramers.

Plausible candidates of pigments for the unique energy-transfer processes in the *Cyanophora* PSI tetramer are the couplings of triply stacked Chls in the monomer1(1’)- monomer2(2’) interface and the Chl-*β*-carotene interactions in the interface between monomer1(1’) and monomer2(2’) and between monomer1(1 ‘) and monomer2’(2) (Supplementary Fig. 13). In particular, the characteristic interactions among Chl823/824/846 (Supplementary Fig. 13) may produce low-energy Chls and characterize time evolution of fluorescence spectra (Fig. 6). In addition, the interactions of Chls-*β*-carotenes, Chl823/824/846-BCR849/BCR852 (Supplementary Fig. 13), may serve as either typical Car → Chl energy transfer^42^ or energy quenching^43,44^. These unique interactions between Chls and *β*-carotenes would be required for the regulation of excitation energy in *Cyanophora*.

### *Cyanophora* PSI tetramers exist *in vivo*

Previous studies showed that *Cyanophora* PSI cores isolated exists in two forms, monomers and tetramers^5,36^. In this study, we recognized that the *Cyanophora* PSI tetramer is significantly labile. Despite the GraFix method, two additional bands of PSI complexes smaller than the PSI tetramers, presumably PSI dimers and monomers, appear in the second-round trehalose gradient centrifugation containing glutaraldehyde (Supplementary Fig. 1A). Thus, the *Cyanophora* PSI tetramers may be readily dissociated into monomers during preparations of the PSI cores^36^, in consistent with the notion that *Cyanophora* PSI is in the middle of transition from cyanobacterial tetramers to eukaryotic monomers.

The weak interactions among the PSI-monomer units in the tetramer raises a question as to whether *Cyanophora* PSI tetramer indeed exists *in vivo*. We tested trehalose gradient centrifugation after solubilizing the thylakoids with a *n*-dodecyl-*β*-D-maltoside (*β*-DDM) concentration as low as 0.1% (w/v) (Supplementary Fig. 21). The results clearly showed the existence of PSI tetramers even with the solubilization of 0.1% *β*-DDM. The ratio of PSI monomer to tetramer is almost not changed between solubilization by 0.1%and 1% *β*-DDM. Since both PSI tetramers and monomers were observed by the BN-PAGE analysis using thylakoid membranes^5^, it is suggested that *Cyanophora* PSI exists in both tetrameric and monomeric forms *in vivo*. Further *in situ* study by cryo-electron tomography will be required for answering this question.

## Conclusions

This study has demonstrated that the eukaryotic PSI from *Cyanophora* can form a tetrameric structure that is remarkably different from the *Anabaena* PSI tetramer. This seems to be caused by both the absence of PsaK in two of the four monomers and the unique structure of PsaL. The tetramer association of *Cyanophora* PSI is rather weak, suggesting that the tetramer can be readily dissociated into monomers. Since other photosynthetic eukaryotes have a monomeric PSI core without PSI tetramers, these structural features imply that *Cyanophora* PSI represents an evolutionary turning-point between cyanobacteria and other photosynthetic eukaryotes.

## Methods

### Purification and characterization of the PSI tetramer from *Cyanophora*

The glaucophyte alga *Cyanophora paradoxa* NIES-547 was grown in 5 L of BG11 medium supplemented with 10 mM Hepes-KOH (pH 8.0) and 5 mL of KW21 (Daiichi Seimo) at a photosynthetic photon flux density of 30 μmol photons m^-2^ s^-1^ at 30°C with bubbling of air containing 3% (v/v) CO2. Note that KW21 is helpful for the growth of photosynthetic organisms as employed for various algae^45–47^. Thylakoid membranes were prepared after disruption of the cells with glass beads as described previously^48^ and suspended in a buffer containing 0.2 M trehalose, 20 mM Mes-NaOH (pH 6.5), 5 mM CaCl_2_, and 10 mM MgCl_2_. The thylakoids were solubilized with 1% (w/v) *β*-DDM at a Chl concentration of 0.25 mg mL^-1^ for 30 min on ice in the dark with gentle stirring. After centrifugation at 20,000 × g for 10 min at 4°C, the resultant supernatant was loaded onto a linear trehalose gradient of 10–40% (w/v) in a medium containing 20 mM Mes-NaOH (pH 6.5), 0.2 M NaCl, and 0.1% *β*-DDM. After centrifugation at 154,000 × g for 18 h at 4°C (P40ST rotor; Hitachi), the PSI-tetramer fraction was obtained in a 25–30% trehalose layer and then concentrated using a 100 kDa cut-off filter (Amicon Ultra; Millipore) at 4,000 × g.

Subunit composition of the PSI was analyzed by a 16–22% SDS-polyacrylamide gel electrophoresis (PAGE) containing 7.5 M urea^49^ (Supplementary Fig. 1B). The sample (2 μg of Chl) was incubated for 10 min at 60°C after addition of 3% lithium lauryl sulfate and 75 mM dithiothreitol. A standard molecular weight marker (SP-0110; APRO Science) was used. The subunit bands separated were identified by mass spectrometry analysis as described previously^50^. An absorption spectrum of the PSI was measured at 77 K using a spectrometer equipped with an integrating sphere unit (V-650/ISVC-747; JASCO)^51^ (Supplementary Fig. 1C). A steady-state fluorescence spectrum of the PSI was recorded at 77 K using a spectrofluorometer (FP-8300/PMU-183; JASCO)^52^ (Supplementary Fig. 1D). Pigment composition of the PSI was analyzed as described in Nagao et al.^53,54^, and the elution profile was monitored at 440 nm (Supplementary Fig. 1E).

### GraFix-treatment of the PSI tetramer

Initial attempts of cryo-grid preparation showed that the PSI tetramers tended to dissociate, and the GraFix method^38^ was therefore used in the last centrifugation step to produce cross-linked samples for cryo-EM analysis in the presence of 0–0.05% glutaraldehyde from top to bottom in the gradient. A fraction of the tetramers was recovered, and a buffer containing 160 mM glycine, 50 mM Mes-NaOH (pH 6.5), 10 mM MgCl_2_, 5 mM CaCl_2_, and 0.03% *β*-DDM was added to stop the cross-linking reaction. The fraction was then concentrated using a 150 kDa cut-off filter (Apollo; Orbital Biosciences, USA) at 4,000 × g, with a buffer exchanged to 50 mM Mes-NaOH (pH 6.5), 10 mM MgCl_2_, 5 mM CaCl_2_, and 0.03% *β*-DDM. The concentrated PSI tetramer was stored in liquid nitrogen until use.

### Cryo-EM data collection

For cryo-EM experiments, 2 μL of sample solution was applied onto a holey carbon grid (Quantifoil R2/1, Cu 300 mesh) covered with a thin amorphous carbon film. The concentrations of uncross-linked and cross-linked samples used are 7 μg and 48 μg of Chl mL^-1^, respectively. The sample-loaded grids were incubated for 30 s in a chamber of an FEI Vitrobot Mark IV at 4°C and 100% humidity. After washing with 2 μL of the solution without trehalose to increase image contrast, the grids were blotted with filter papers for 5 s and then immediately plunge-frozen into liquid ethane. The grids were examined with a 200 kV cryo-electron microscope (Talos Arctica; Thermo Fisher Scientific) incorporating a field emission gun and a direct electron detector (Falcon 3EC; Thermo Fisher Scientific). The conditions for the cryo-EM data collection are summarized in Supplementary Table 1.

### Cryo-EM image processing

Cryo-EM movies were recorded at a nominal magnification of × 92,000 using the Falcon 3EC detector in a linear mode (calibrated pixel size of 1.093 Å). The movie frames were aligned and summed using the MotionCor2 software^55^, and the contrast transfer function (CTF) was estimated using the Gctf program^56^. The 3D structures are reconstructed using RELION^57^.

For structural analysis of the unfixed (uncross-linked) sample, in total 1,603,082 particles were automatically picked from 4,515 micrographs, and they are subjected to reference-free 2D classification. The procedure of the structural analysis is summarized in Supplementary Fig. 2. Tetramers and dimers were observed in the 2D classification. To determine a structure in the tetrameric form (a dimer of dimers), particles in classes viewed from the top or bottom (perpendicular to the membrane) containing only tetramers and classes viewed from the side (parallel to the membrane) probably containing tetramers and dimers were selected (1,010,216 particles) and subjected to three rounds of 3D classification with a C2 symmetry. The reference model used in the 3D classification was generated in RELION. After removing bad particles in the low-resolution, good particles were further subjected to second-round 2D classification. Finally, 145,567 particles were selected and used for the 3D reconstruction. The final map in the tetrameric form was reconstructed with a C2 symmetry at 4.0 Å resolution, which was estimated by the gold standard Fourier shell correlation at 0.143 criterion^58^. To improve the resolution in each monomeric unit, we performed focused 3D classifications using masks covering each monomeric unit. To determine higher-resolution structures in each monomeric unit (monomer1 and monomer2), we combined particles in the dimeric and tetrameric forms. Dimeric particles after the first 2D classification were selected (622,323 particles) and subjected to first-round 3D classification for the dimers. Particles in good classes were selected (416,757 particles) and combined with tetrameric particles (145,567 particles). As the tetrameric particle is a dimer of dimers and has a two-fold symmetry, the particle orientation was expanded with a C2 symmetry before joining the particle (291,134 particle orientations). The joined particle set was subjected to second-round 3D classification using a mask covering a dimer. Particles in good classes were selected (660,237 particle orientations) and subjected to two rounds of the focused classification using masks covering each monomeric unit. The final maps of monomer1 and monomer2 were reconstructed from 70,920 and 110,380 particles at 3.3 Å and 3.2 Å resolutions, respectively (Supplementary Fig. 3). Local resolutions were estimated using RELION (Supplementary Fig. 3).

For structural analysis of the GraFix-treated sample, in total 724,316 particles were automatically picked from 3,205 micrographs and then used for reference-free 2D classification. For structure of the GraFix PSI tetramer, in total 426,961 particles were selected from good 2D classes and subsequently subjected to two rounds of the 3D classification with or without the C2 symmetry. The initial model used for the first 3D classification was the structure of unfixed PSI tetramer at 4.0 Å resolution. As shown in Supplementary Fig. 7C, the structure of the GraFix PSI tetramer was reconstructed from 40,679 particles at an overall resolution of 3.8 Å (Supplementary Fig. 8). Local resolution was estimated using RELION (Supplementary Fig. 8).

### Model building and refinement

The 3.3-Å and 3.2-Å cryo-EM maps were used for the model building of the wild-type PSI monomer1 and monomer2 within the PSI tetramer, respectively. For the PSI-core model building, homology models constructed using Phyre2^59^ were first manually fitted into each map with UCSF Chimera^60^, and then inspected and adjusted individually with Coot^61^. The wild-type PSI monomer1 and monomer2 structures were then refined with phenix.real_space_refine^62^ with geometric restraints for protein-cofactor coordination. For structural analysis of the GraFix PSI tetramer, the wild-type PSI monomer1 and monomer2 within the tetramer were manually fitted into the 3.8-Å cryo-EM map using UCSF Chimera and were then refined with phenix.real_space_refine with geometric restraints for protein-cofactor coordination. The final models were further validated with MolProbity^63^ and EMringer^64^. The statistics for all data collection and structure refinement are summarized in Supplementary Table 1.

### TRF measurement

TRF spectra were recorded by a time-correlated single-photon counting system with a wavelength interval of 1 nm and a time interval of 2.44 ps^65^. A picosecond pulse diode laser (PiL044X; Advanced Laser Diode Systems) was used as an excitation source, and it was operated at 445 nm with a repetition rate of 3 MHz. The details of the TRF-measurement conditions were described previously^66^.

### Phylogenetic analyses of the PsaA, PsaB, PsaK, and PsaL

Alignment and phylogenetic reconstructions were performed using the function “build” of ETE3 v3.1.1^67^ as implemented on the GenomeNet (https://www.genome.jp/tools/ete/). The tree was constructed using FastTree v2.1.8 with the default parameters^68^. The species used for the analysis are *Cyanophora paradoxa*, *Thermosynechococcus elongatus* BP-1, *Synechocystis* sp. PCC 6803, *Anabaena* sp. PCC 7120, *Chaetoceros gracilis*, *Chlamydomonas reinhardtii*, *Cyanidioschyzon merolae*, *Pisum sativum*, and *Zea mays*.

## Data availability

Atomic coordinates and cryo-EM maps for the reported structures of monomer1, monomer2, and the GraFix-treated PSI tetramer have been deposited in the Protein Data Bank under an accession codes 7DR0, 7DR1, and 7DR2, and in the Electron Microscopy Data Bank under the accession codes EMD-30820, EMD-30821, and EMD-30823, respectively. The raw electron microscopy images used to build the 3D structure are available from the corresponding authors upon request.

## Acknowledgements

We thank Drs. Naruhiko Adachi and Masato Kawasaki for helpful assistance during the cryo-EM study. This work was supported by the Platform Project for Supporting Drug Discovery and Life Science Research (Basis for Supporting Innovative Drug Discovery and Life Science Research (BINDS)) from AMED, JSPS KAKENHI grant Nos. JP20H02914 (K.K.), JP20K06528, JP21K19085 (R.N.), JP18J10095 (Y.U.), JP19K22396, JP20H03194 (F.A.), JP16H06553 (S.A.), JP20H05087 (N.M.), and JP17H06433 (J.-R.S.), Takeda Science Foundation (K.K.), and TIA-Kakehashi grant No. TK19-048 (N.M.).

## Author Contributions

R.N. and J.-R.S. conceived the project; K.K., R.N., and T.-Y.J purified the PSI cores and performed biochemical characterizations; Y.U., M.Y., and S.A. performed spectroscopic measurements and analyzed the data; T.S. and N.D. identified PSI subunits by MS analyses; K.K. performed phylogenetic analyses; F.A. and N.M. collected cryo-EM images; K.K. and N.M. processed the EM data and reconstructed the final EM maps; K.K. built the structure model and refined the final models; K.K., R.N., N.M., S.A., and J.-R.S. wrote a draft manuscript; and R.N. and J.-R.S. wrote the final manuscript, and all of the authors joined the discussion of the results.

## Competing interests

Authors declare no competing interests.

## Additional information

Supplementary information is available for this paper at

## Notes

### Competing Interest Statement

The authors have declared no competing interest.

