## Supplementary Information for "Structural insights into an evolutionary turning-point of photosystem I from prokaryotes to eukaryotes"

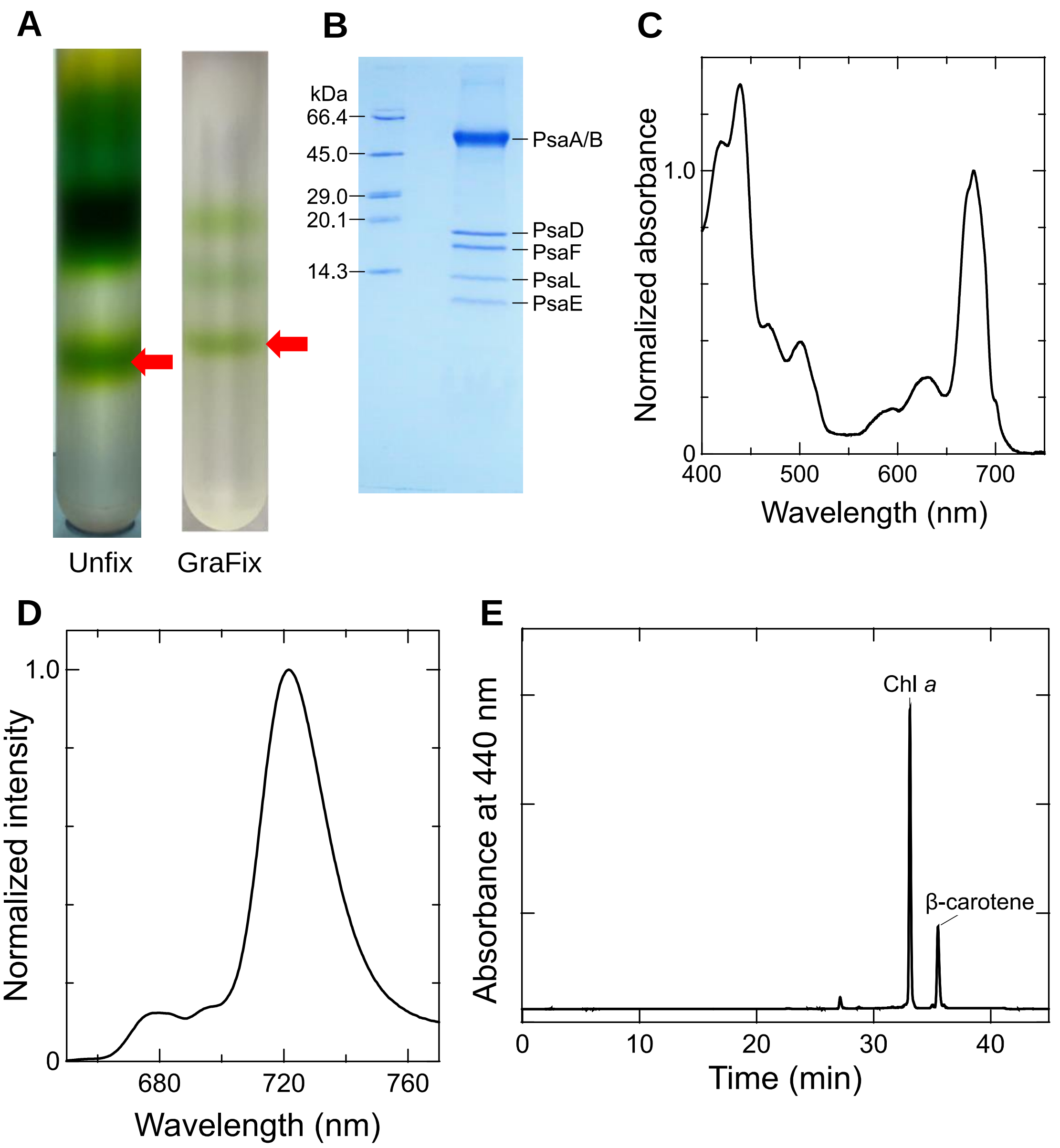

**Supplementary Figure 1. Biochemical and spectroscopic characterizations of the PSI tetramer.**

**A,** Trehalose gradient centrifugation of the unfix (left) and GraFix (right) PSI preparation from *Cyanophora paradoxa.* Red arrows indicate PSI tetramer. **B,** SDS-PAGE analysis of the PSI tetramer obtained from trehalose gradient centrifugation shown in panel **A**. **C,** Absorption spectrum of the PSI tetramer measured at 77 K. **D,** Normalized fluorescence spectrum of the PSI tetramer excited at 445 nm and measured at 77 K. **E,** HPLC analysis of pigments extracted from the PSI tetramer monitored at 440 nm.

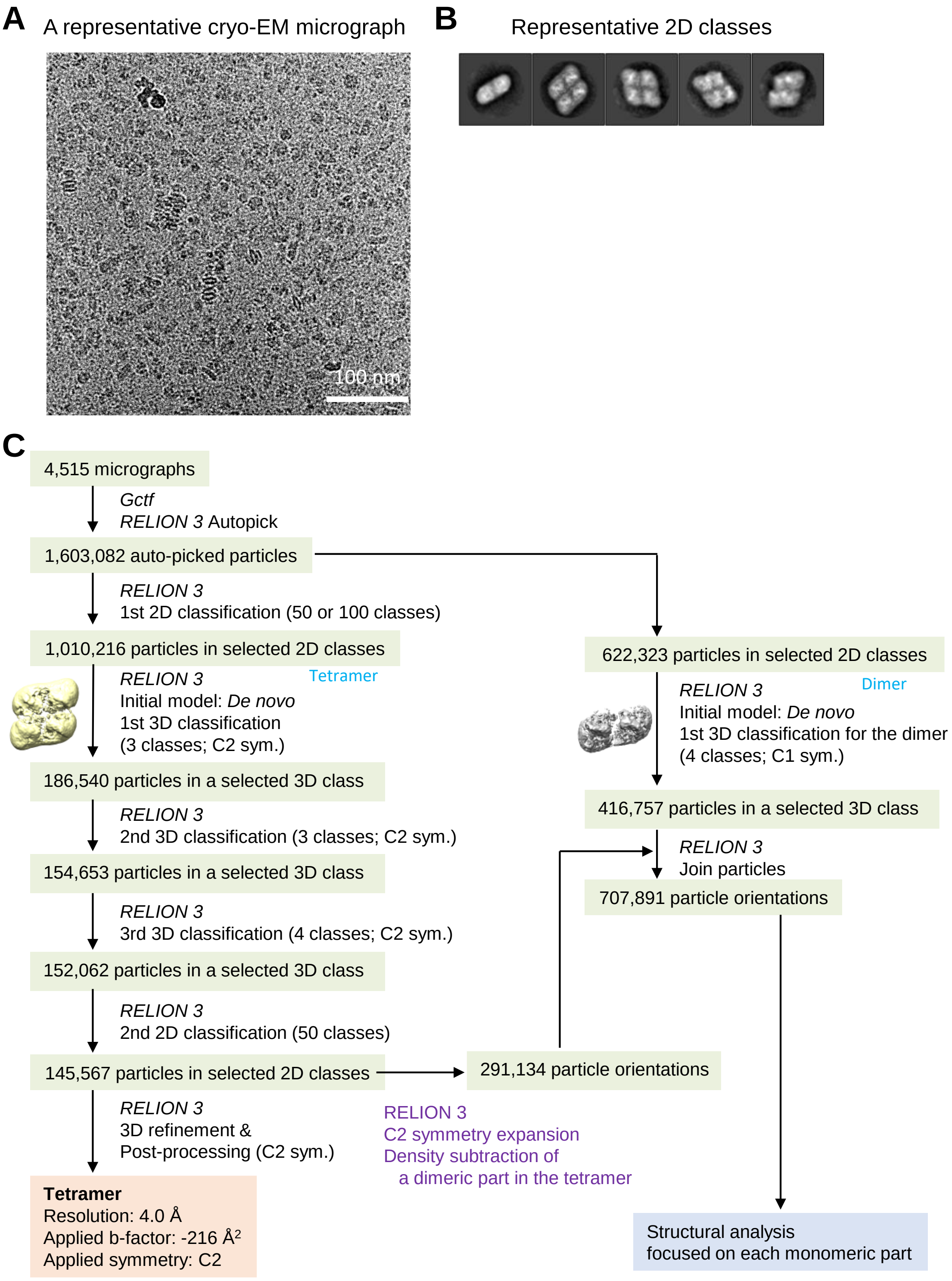
A
A representative cryo-EM micrograph
100 nm
B
Representative 2D classes
C
4,515 micrographs
Gctf
RELION 3 Autopick
1,603,082 auto-picked particles
RELION 3
1st 2D classification (50 or 100 classes)
1,010,216 particles in selected 2D classes
RELION 3
Tetramer
Initial model: De novo
1st 3D classification
(3 classes; C2 sym.)
186,540 particles in a selected 3D class
RELION 3
2nd 3D classification (3 classes; C2 sym.)
154,653 particles in a selected 3D class
RELION 3
3rd 3D classification (4 classes; C2 sym.)
152,062 particles in a selected 3D class
RELION 3
2nd 2D classification (50 classes)
145,567 particles in selected 2D classes
RELION 3
3D refinement &
Post-processing (C2 sym.)
Tetramer
Resolution: 4.0 Å
Applied b-factor: -216 Å2
Applied symmetry: C2
622,323 particles in selected 2D classes
RELION 3
Dimer
Initial model: De novo
1st 3D classification for the dimer
(4 classes; C1 sym.)
416,757 particles in a selected 3D class
RELION 3
Join particles
707,891 particle orientations
291,134 particle orientations
RELION 3
C2 symmetry expansion
Density subtraction of
a dimeric part in the tetramer
Structural analysis
focused on each monomeric part

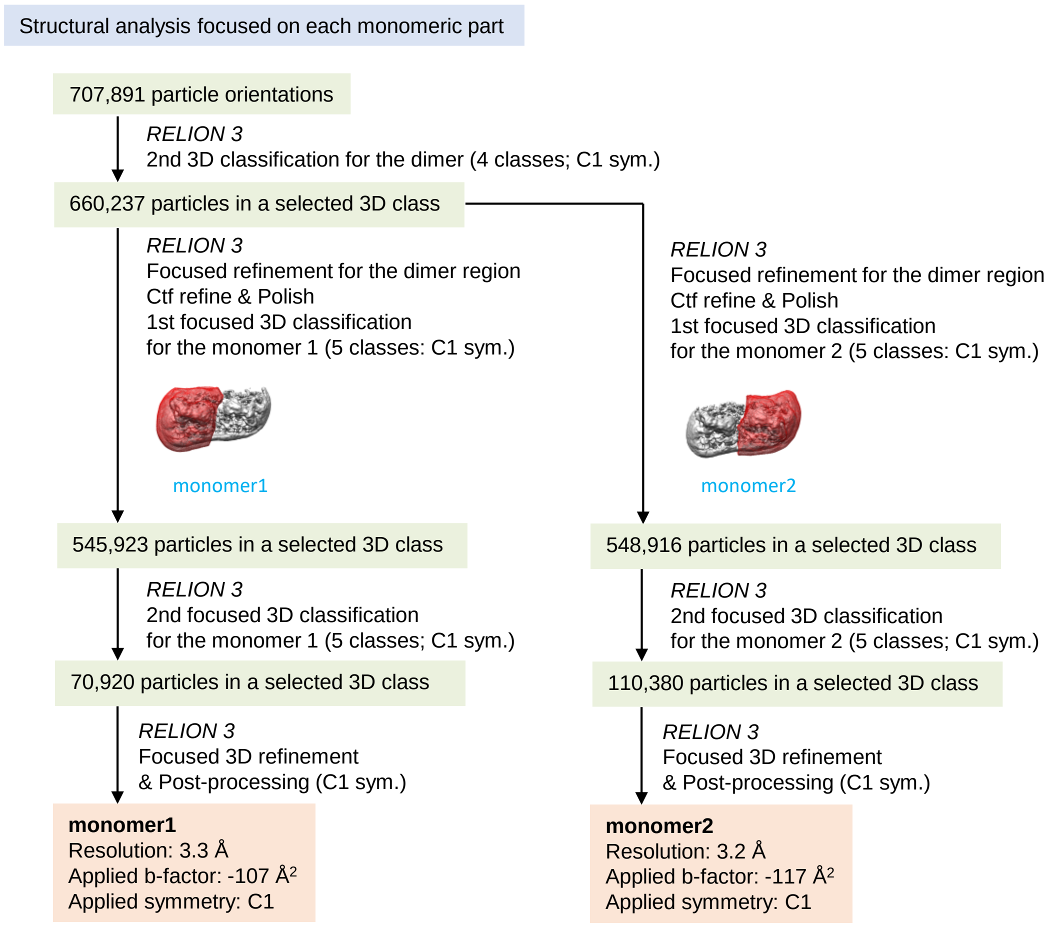

**Supplementary Figure 2. Cryo-EM data collection and processing of the wild-type PSI tetramer.**

**A,** A representative cryo-EM micrograph of the PSI tetramer. **B,** Representative 2D classes of the PSI tetramer. The box size is 437 Å. **C,** A schematic flowchart showing the classification scheme for the PSI tetramer. The overall PSI-tetramer structure was reconstructed at 4.0 Å resolution from 145,567 particles, whereas monomer1 and monomer2 structures from the subtracted particles were reconstructed at resolutions of 3.3 and 3.2 Å, respectively. See Methods section for more details.

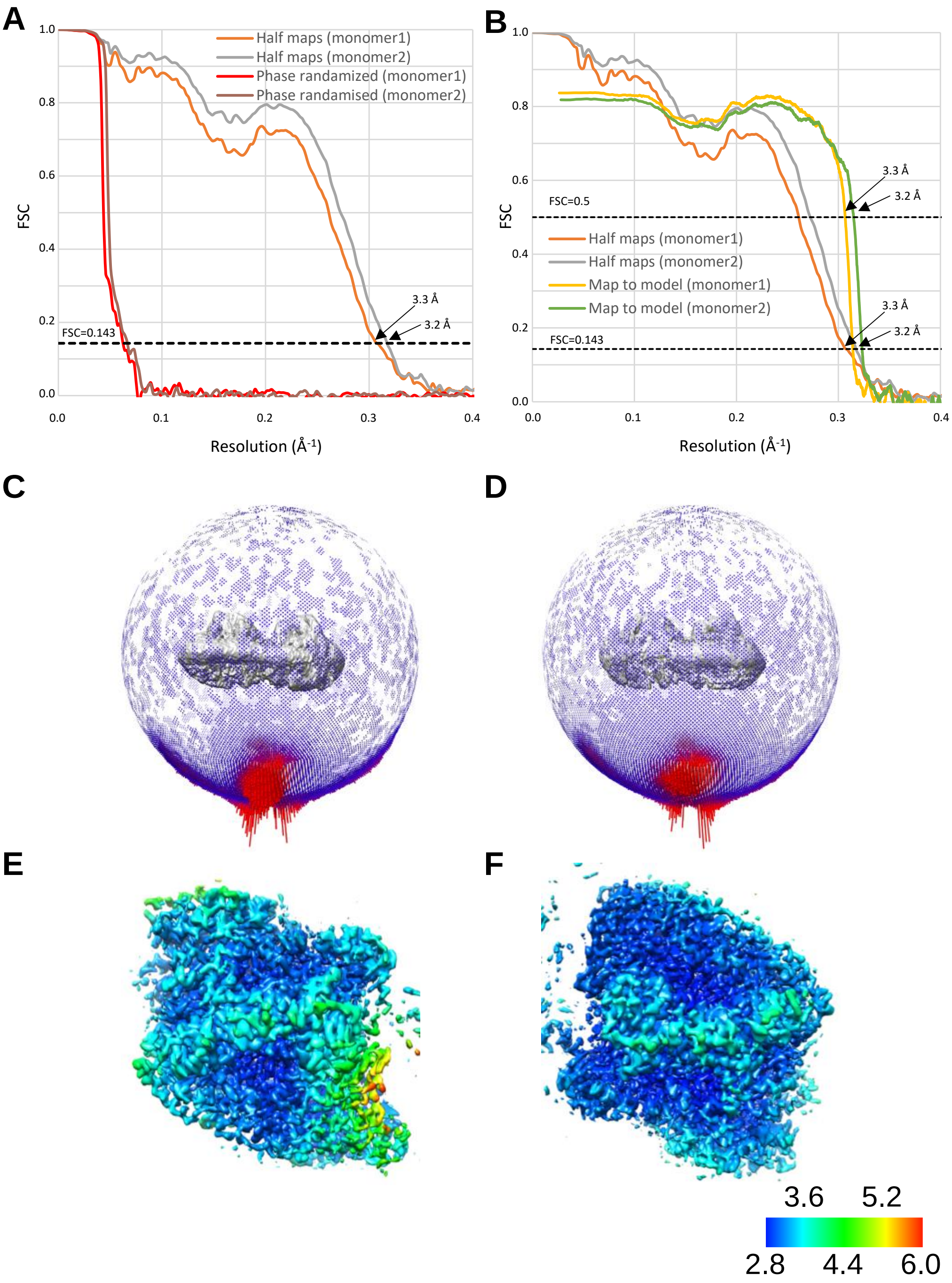

**Supplementary Figure 3. Evaluation of the cryo-EM map quality.**
**A,** FSC curves of the wild-type monomer1 and monomer2 for independently refined half-maps (half-maps) and phase randomized maps. **B,** FSC curves of the wild-type monomer1 and monomer2 for independently refined half-maps (half-maps) and full map vs. model (map to model). **C, D,** Angular distributions of the particles used for reconstruction of monomer1 (**C**) and monomer2 (**D**). Each cylinder represents one view, and the height of the cylinder is proportional to the number of particles for that view. **E, F,** Local resolution maps of monomer1 (**E**) and monomer2 (**F**).

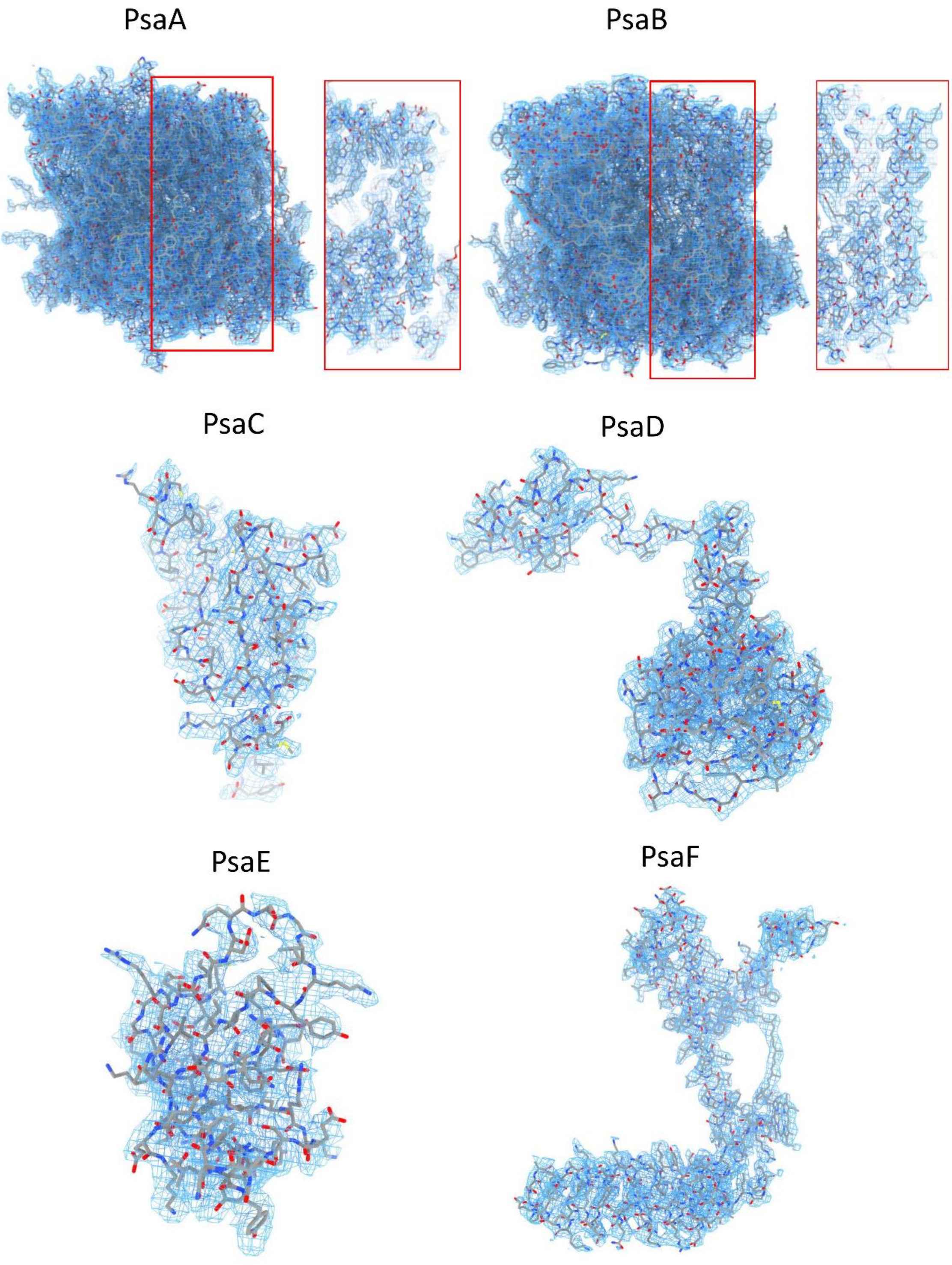
PsaA
PsaB
PsaC
PsaD
PsaE
PsaF

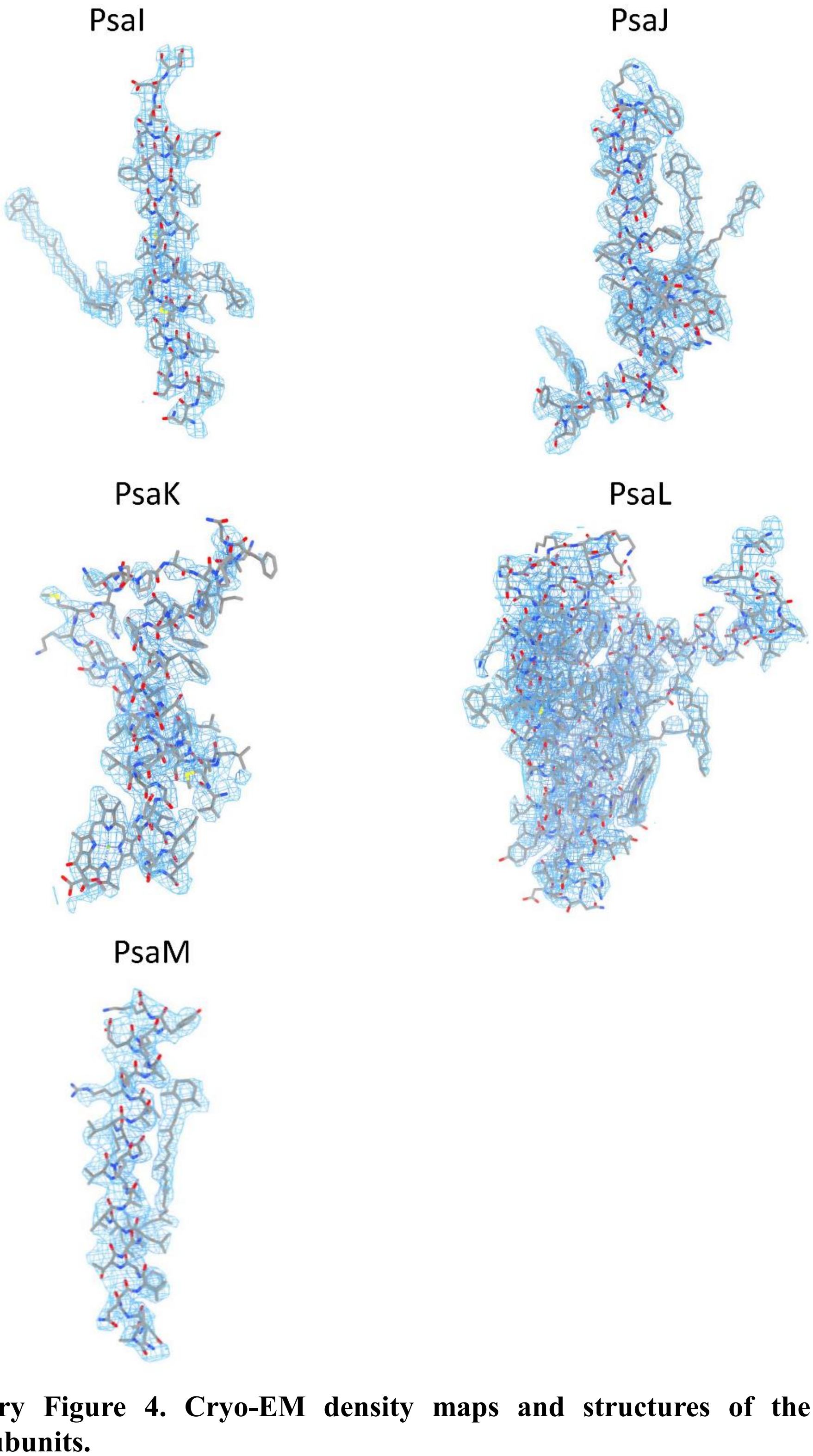

**Supplementary Figure 4. Cryo-EM density maps and structures of the PSI-monomer1 subunits.**

The densities for each subunit of PSI are shown as blue meshes, and the corresponding models are shown as gray sticks. Red boxes indicate the enlarged views for a part of the individual PsaA and PsaB subunits.

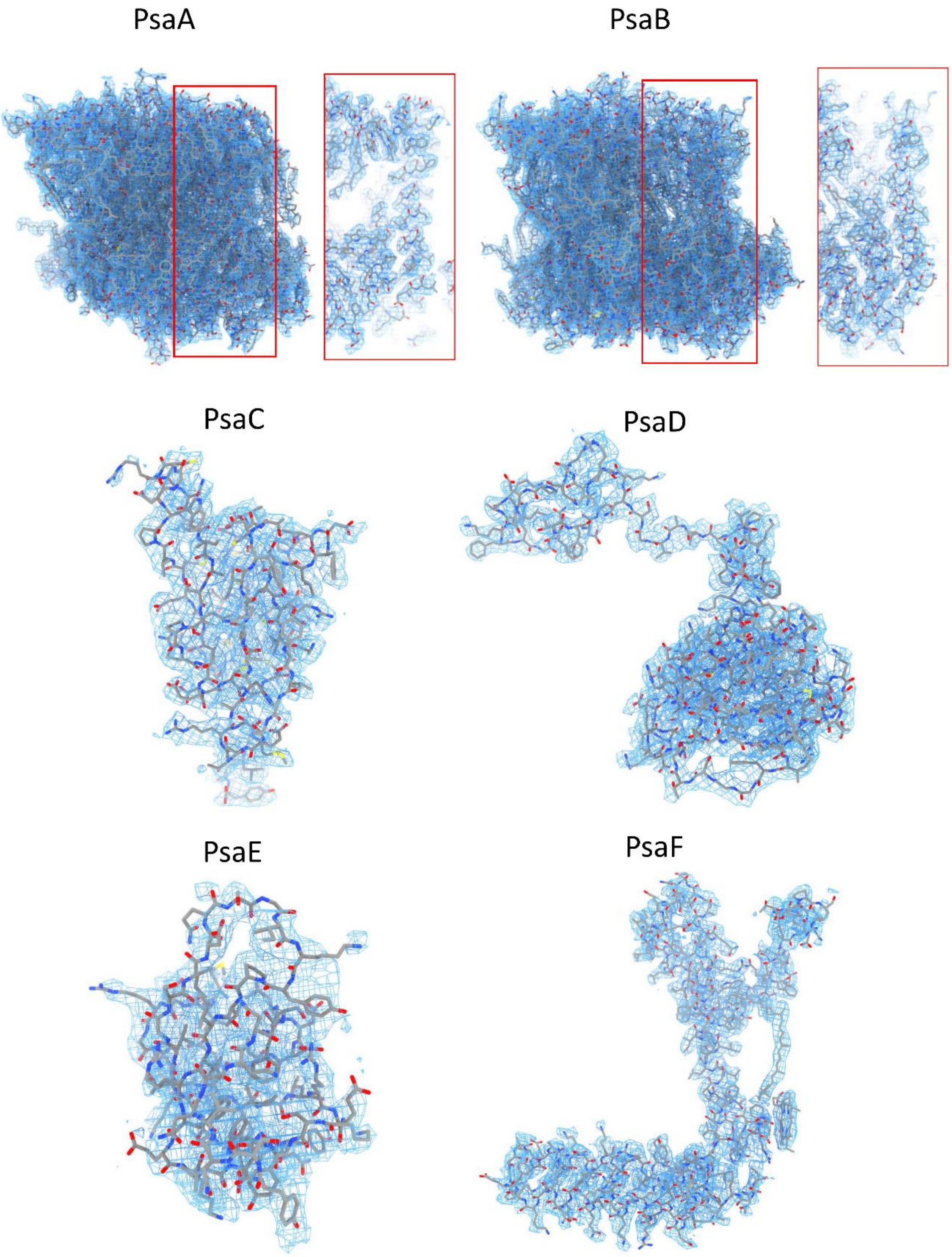
PsaA
PsaB
PsaC
PsaD
PsaE
PsaF

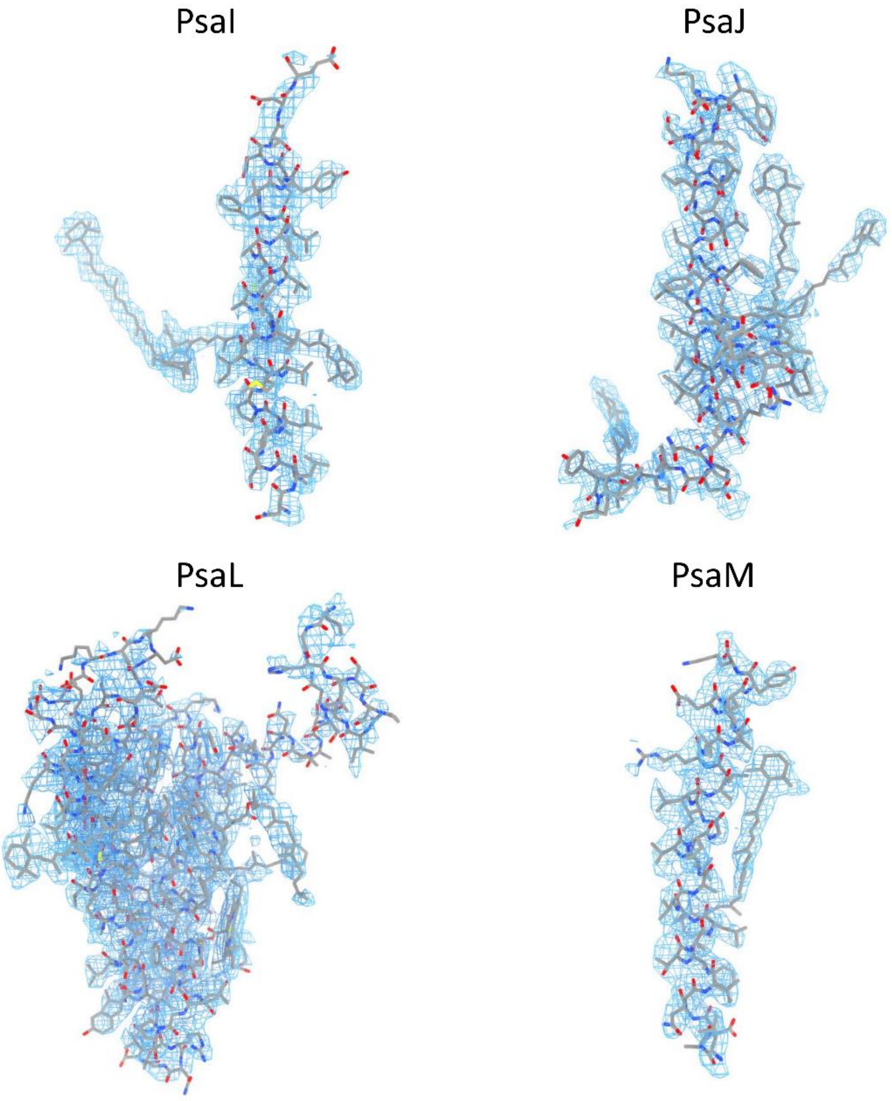

**Supplementary Figure 5. Cryo-EM density maps and structures of the PSI-monomer2 subunits.**

The densities for each subunit of PSI are shown as blue meshes, and the corresponding models are shown as gray sticks. Red boxes indicate the enlarged views for a part of the individual PsaA and PsaB subunits.

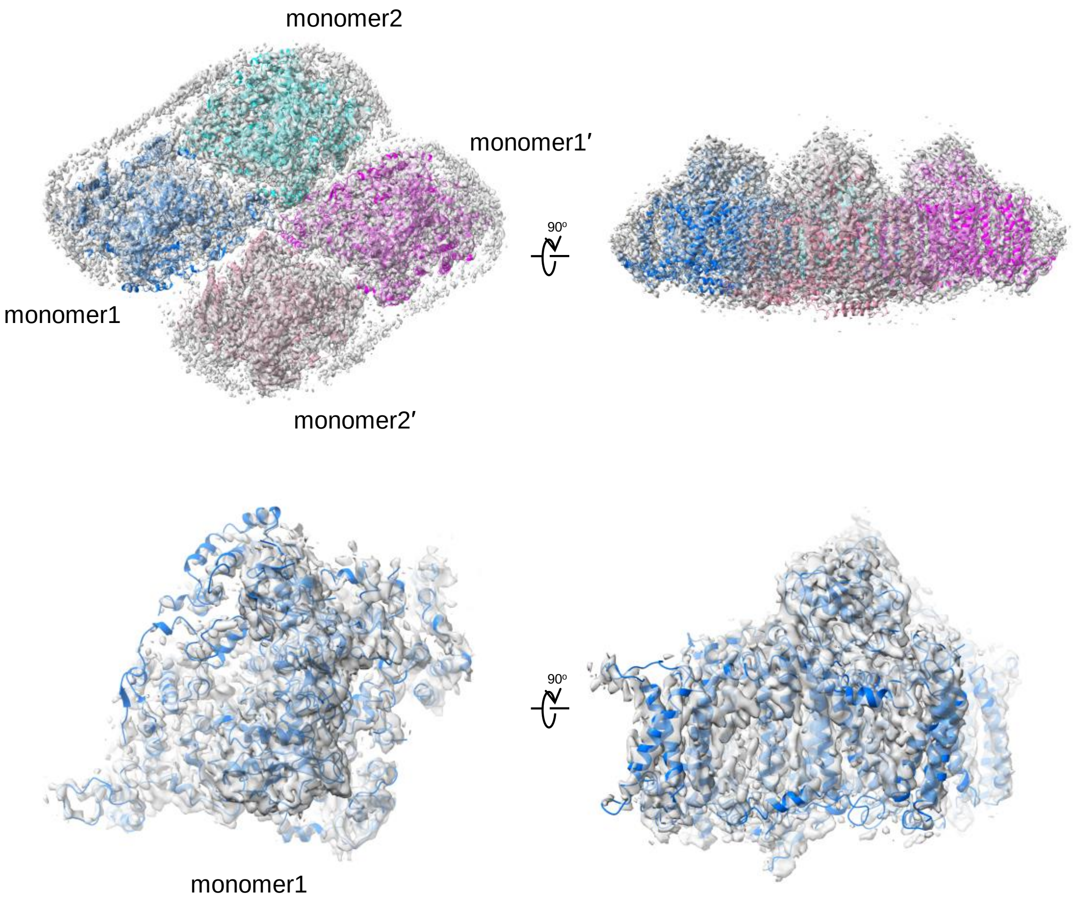

**Supplementary Figure 6. Overall structure of the wild-type PSI tetramer.**
The 3D cryo-EM density maps of the PSI tetramer superimposed with a cartoon model of the PSI tetramer, with a view along the membrane normal from the stromal side (upper left) and its side view (upper right). Lower panel shows the superimposition of the monomer1 structure with its cryo-EM density.

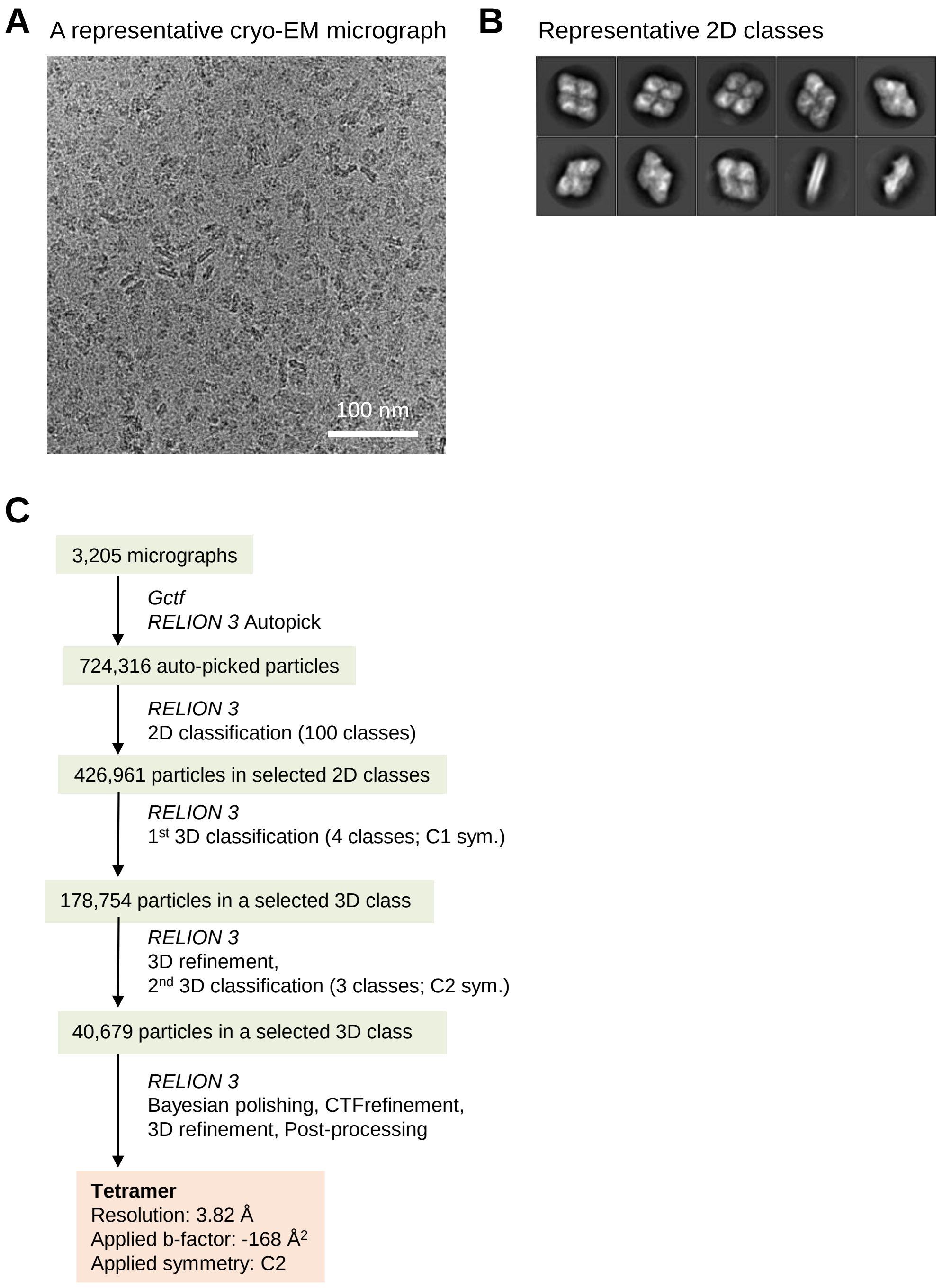

**Supplementary Figure 7. Cryo-EM data collection and processing of the GraFix PSI tetramer.**

**A,** A representative cryo-EM micrograph of the PSI tetramer. **B,** Representative 2D classes of the PSI tetramer. The box size is 437 Å. **C,** A schematic flowchart showing the classification scheme for the PSI tetramer. The overall PSI tetramer structure was reconstructed at 3.8 Å resolution from 40,679 particles. See Methods section for more details.

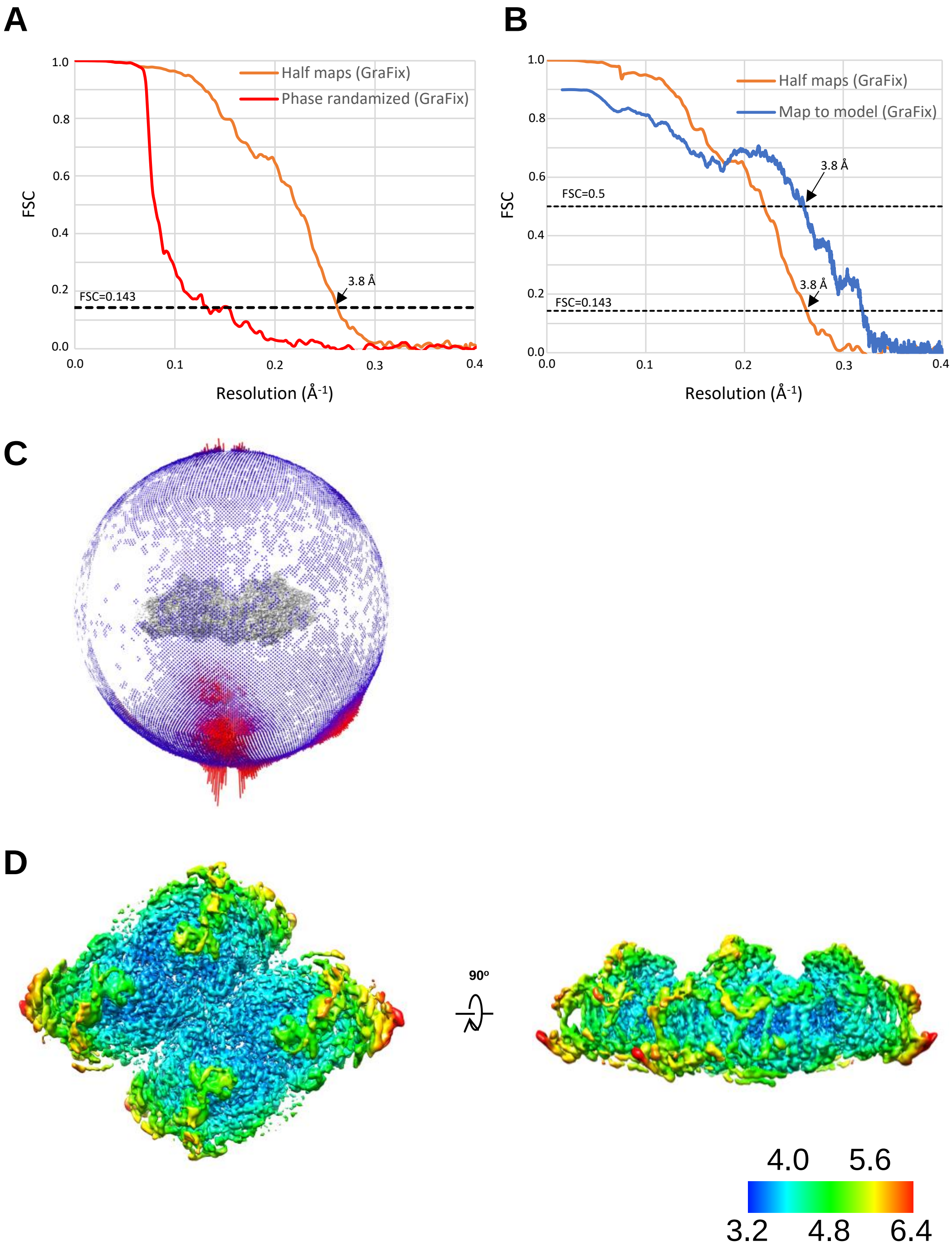

**Supplementary Figure 8. Evaluation of the cryo-EM map quality.**

**A,** FSC curves of the GraFix PSI tetramer for independently refined half-maps (half-maps) and phase randomized maps. **B,** FSC curves of the GraFix PSI tetramer for independently refined half-maps (half-maps) and full map vs. model (map to model). **C,** Angular distribution of the particles used for reconstruction of the GraFix PSI tetramer. Each cylinder represents one view, and the height of the cylinder is proportional to the number of particles for that view. **D,** Local resolution map of the GraFix PSI tetramer.

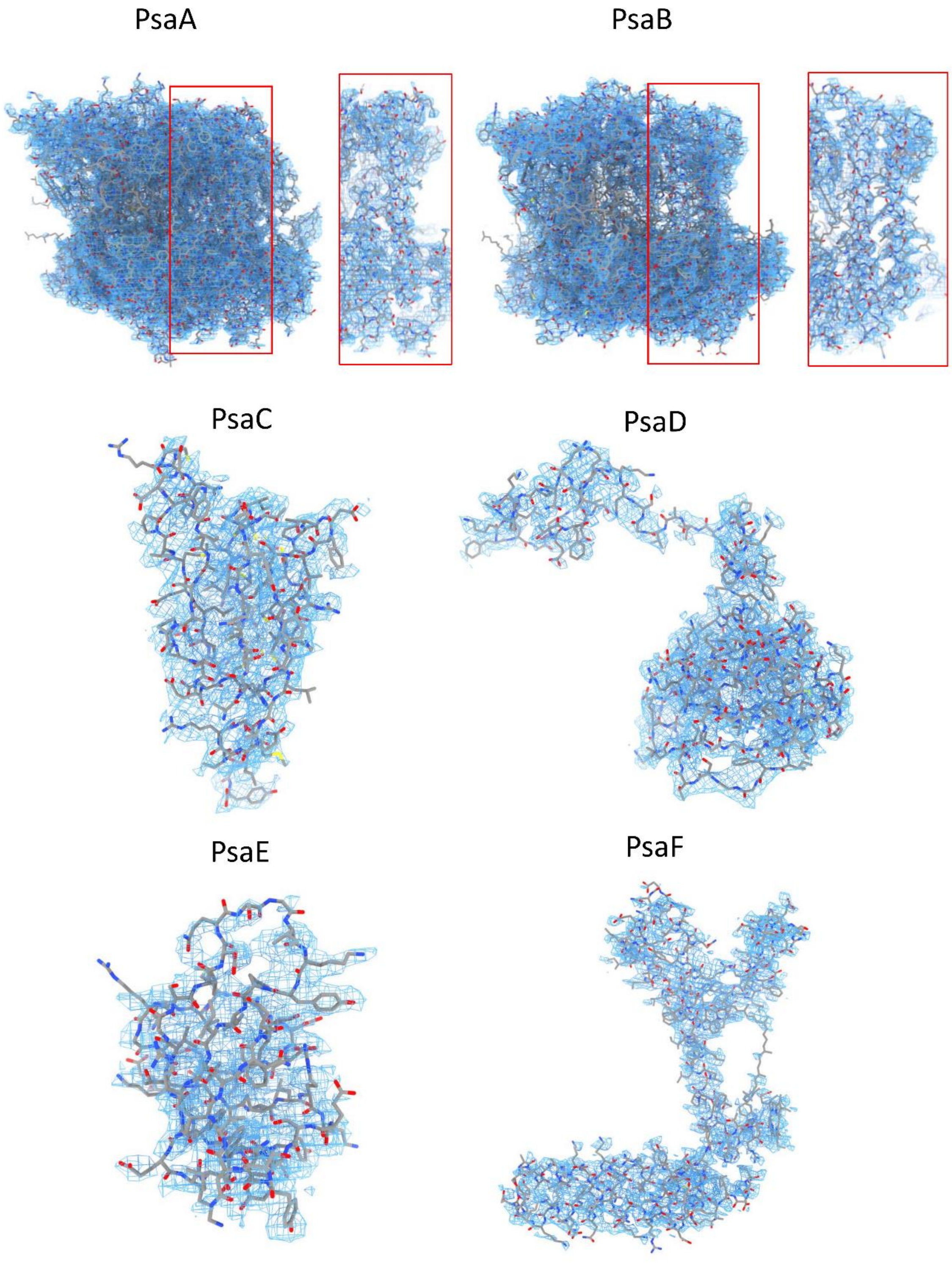
PsaA
PsaB
PsaC
PsaD
PsaE
PsaF

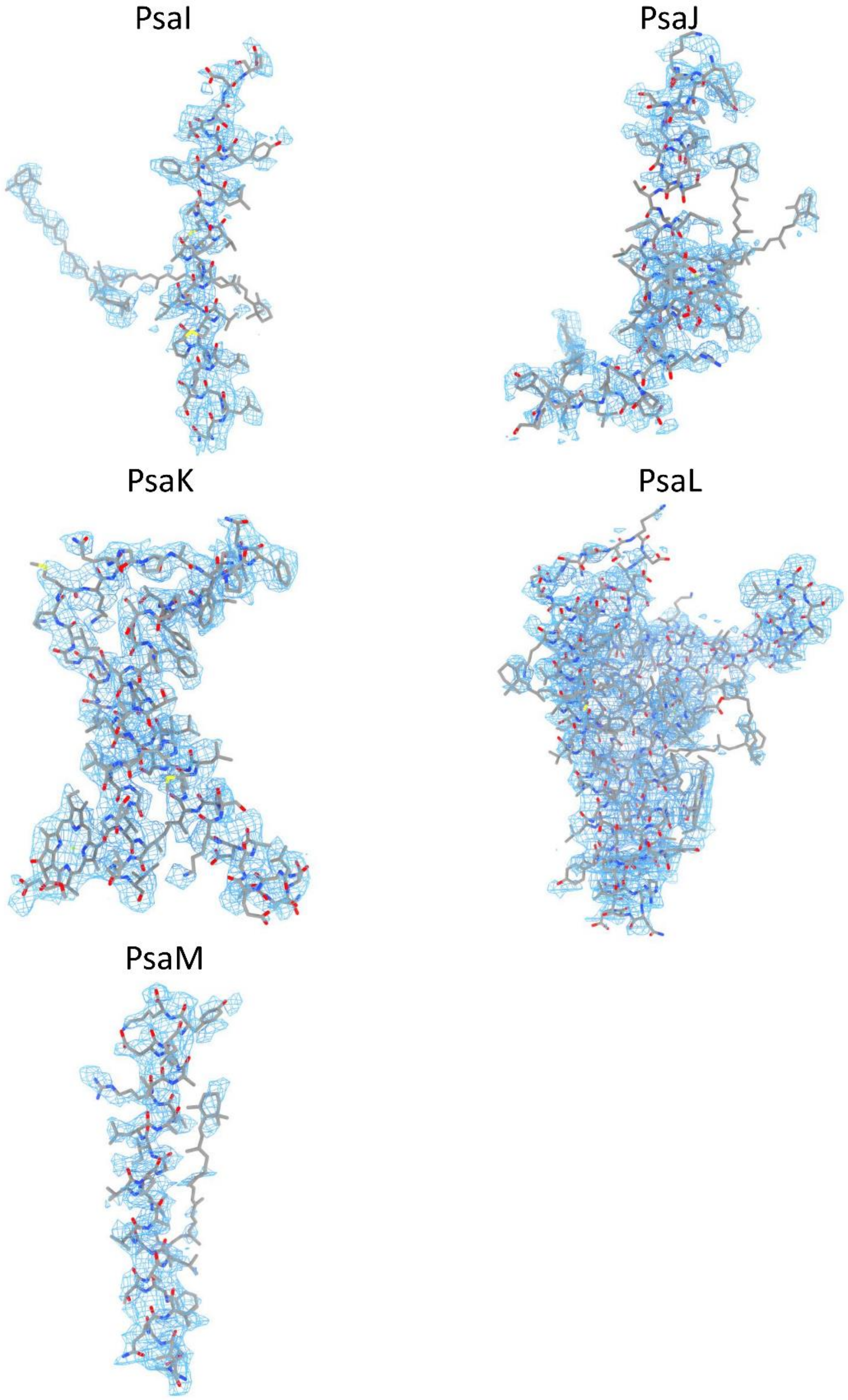

**Supplementary Figure 9. Cryo-EM density maps and structures of the GraFix PSI tetramer subunits.**

The densities for each subunit of PSI are shown as blue meshes, and the corresponding models are shown as gray sticks. Red boxes indicate the enlarged views for a part of the individual PsaA and PsaB subunits.

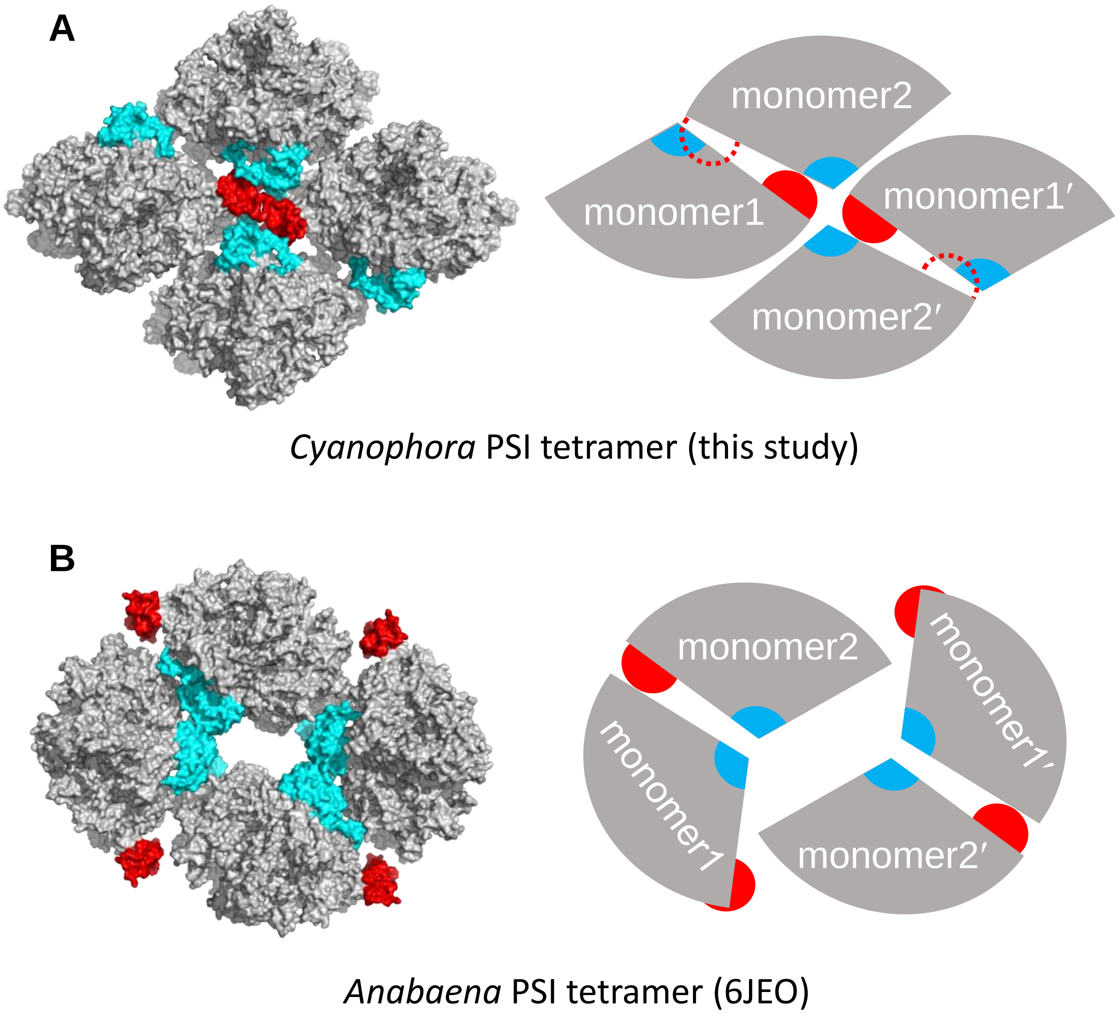

**Supplementary Figure 10. Comparison of the organization patterns of the PSI tetramer between *Cyanophora* and *Anabaena*.**

**A,** *Cyanophora* PSI tetramer. **B,** *Anabaena* PSI tetramer (PDB: 6JEO). Red and cyan represent PsaK and PsaL, respectively. The areas cycled by red dashed lines in right side of panel **A** represent the sites of PsaK that are in conflict with the adjacent monomer and lost in the structure.

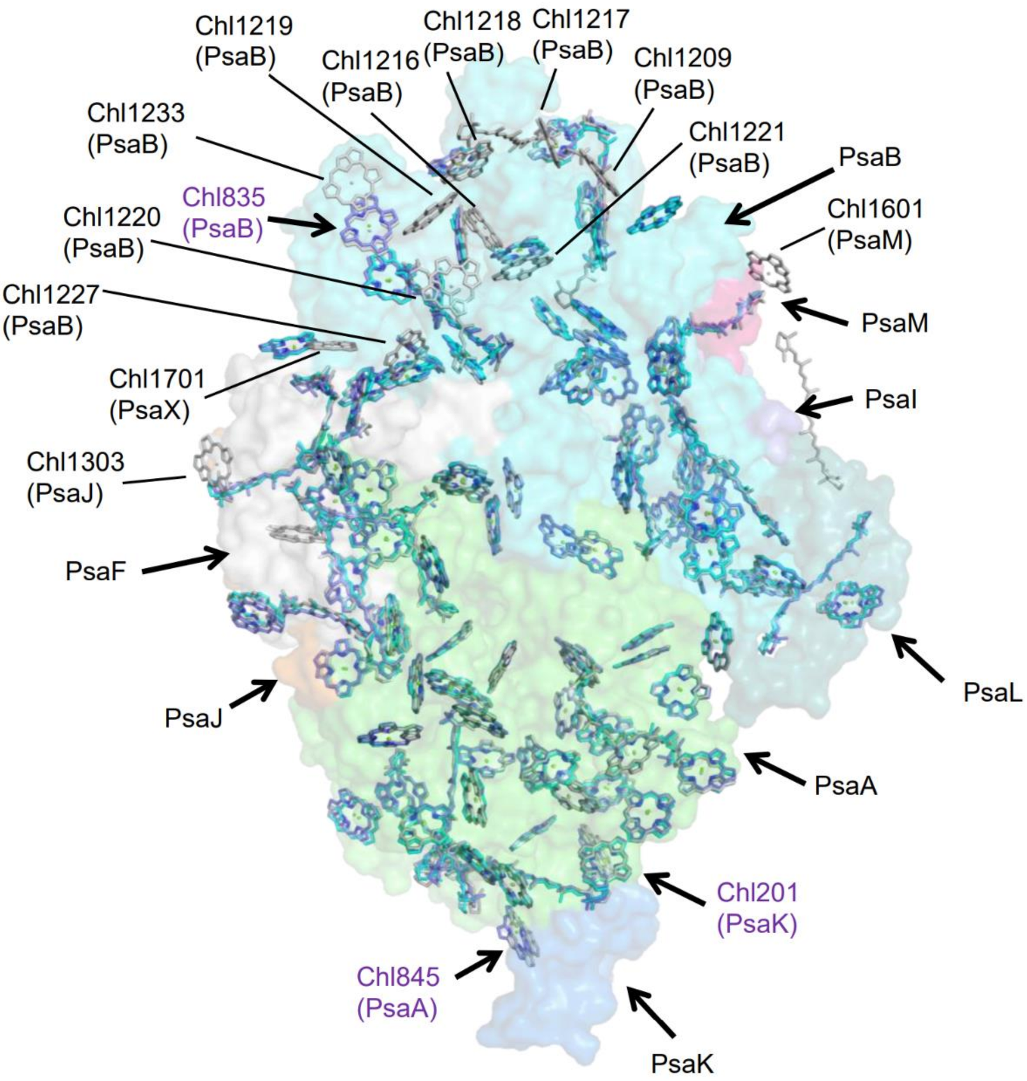

**Supplementary Figure 11. Superposition of the pigment molecules of the PSI tetramer with those of the PSI trimer.**

Arrangement of the pigments in monomer1 (purple), monomer2 (cyan), and a monomer unit of the PSI trimer (PDB: 1JB0) (gray), with protein surface model viewed along the membrane normal from the stromal side. The stromal side extrinsic subunits PsaC, PsaD, and PsaE are deleted in the model for clarity.

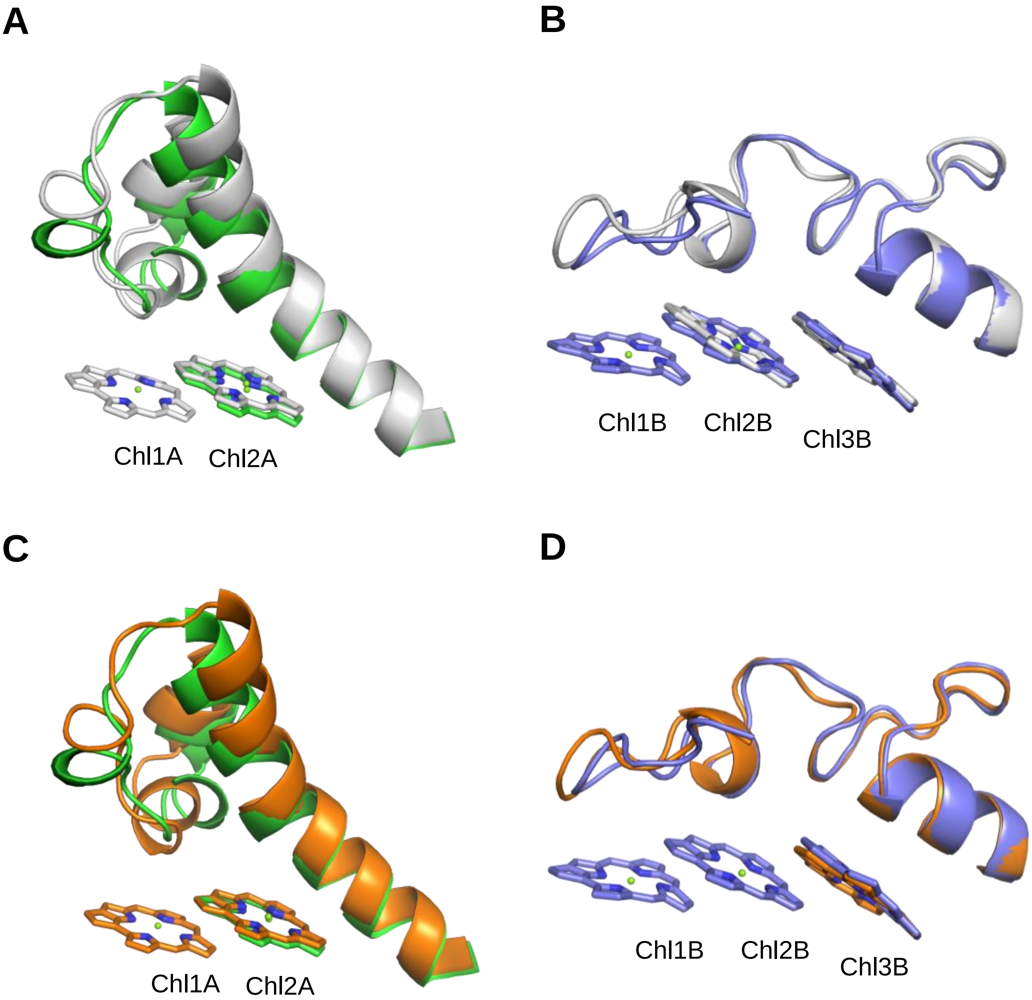

**Supplementary Figure 12. Comparison of the Low1 and Low2 sites between the *Cyanophora* PSI and other cyanobacterial PSI.**

**A,** Superposition of the Low1 site in the *Cyanophora* PSI monomer1 (gray) and the PSI-monomer unit of *Gloeobacter violaceus* PCC 7421 (PDB: 7F4V) (green). **B,** Superposition of the Low2 site in the *Cyanophora* PSI monomer1 (gray) and the PSI-monomer unit of *Thermosynechococcus vulcanus* NIES-2134 (PDB: 6K33) (blue). **C,** Superposition of the Low1 site in the *Cyanophora* PSI monomer2 (orange) and the PSI-monomer unit of *Gloeobacter violaceus* PCC 7421 (PDB: 7F4V) (green). **D,** Superposition of the Low2 site in the *Cyanophora* PSI monomer2 (orange) and the PSI-monomer unit of *Thermosynechococcus vulcanus* NIES-2134 (PDB: 6K33) (blue).

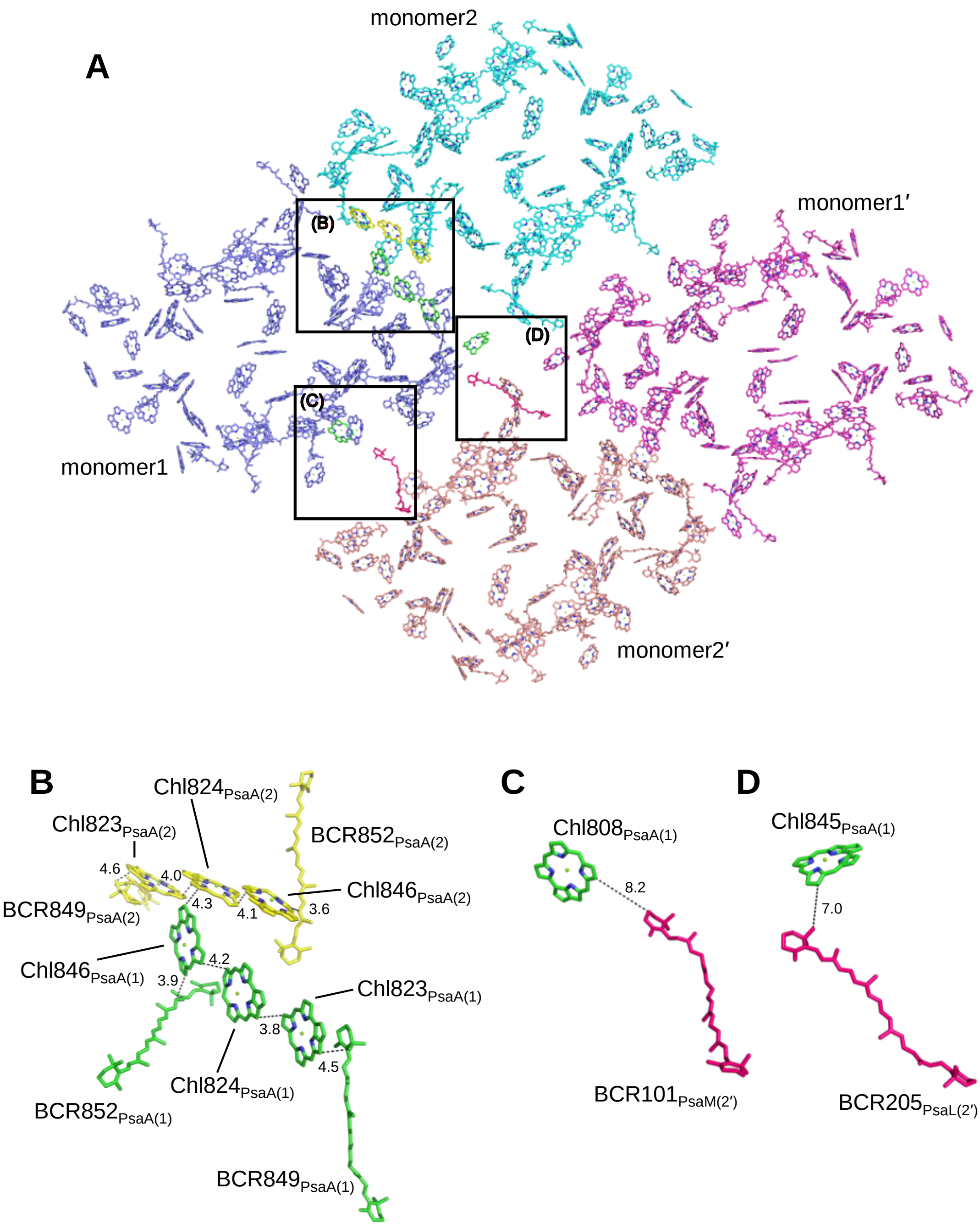

**Supplementary Figure 13. Arrangement of pigments in the *Cyanophora* PSI tetramer and their interactions.**

**A,** Arrangement of the pigments in the PSI tetramer viewed along the membrane normal from the stromal side. Pigments of monomer1, monomer2, monomer1′, and monomer2′ are colored in blue, cyan, magenta, and pink, respectively. A triply stacked Chl cluster in monomer1, and a triple Chl cluster in monomer2 that interacts with monomer1, are colored in green and yellow, respectively. Boxed areas are enlarged in the following panels. **B,** Interactions of pigments between monomer1 (green) and monomer2 (yellow) shown in box (**B**) of panel **A**. **C, D,** Interactions of two Chls in monomer1 and two *β*-carotenes in monomer2′ shown in boxes (**C**) and (**D**) of panel **A**.

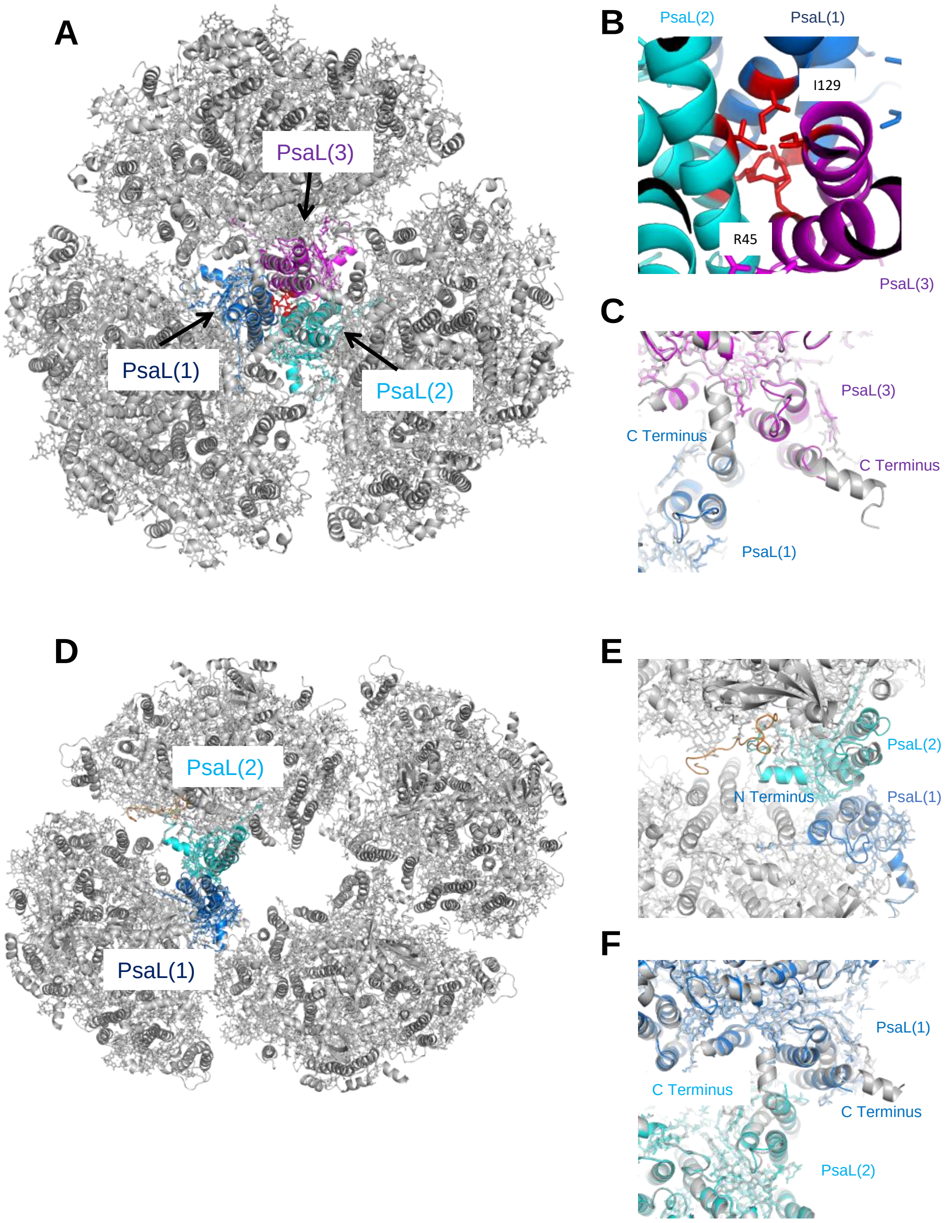

**Supplementary Figure 14. Structural modeling of *Cyanophora* PSI trimer and tetramer.**

**A,** Superposition of a *Cyanophora* PSI monomer with the trimeric structure of PSI from *T. elongatus* (PDB: 1JB0), viewed along the membrane normal from the stromal side. The region having steric hindrances is colored in red. PsaL in monomer1, monomer2, and monomer3 are depicted in blue, cyan, and purple, respectively. **B,** Close-up view of the steric hindrance of PsaL from the lumenal side. **C,** Close-up view of the C-terminus of PsaL from the lumenal side. **D,** Superposition of a *Cyanophora* PSI monomer with the tetrameric structure of PSI from *Anabaena* sp. PCC 7120 (PDB: 6JEO), viewed along the membrane normal from the stromal side. **E,** Close-up view of the N-terminus of PsaL from the stromal side. A loop structure (orange) stands for the N-teminal loop of PsaL in

*Anabaena* sp. PCC 7120. **F,** Close-up view of the C-terminus of PsaL from the lumenal side.

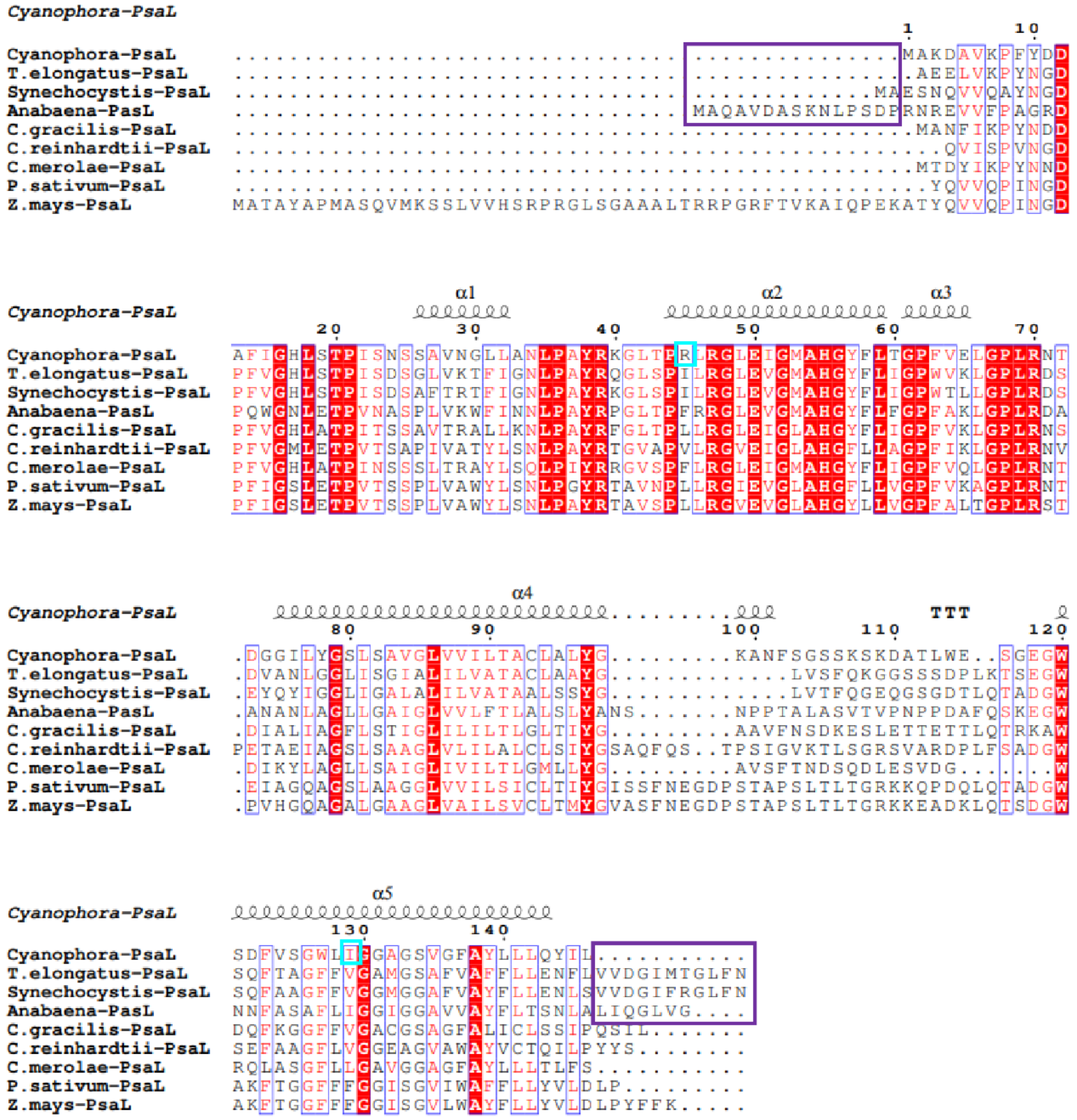

**Supplementary Figure 15. Multiple sequence alignment (CLUSTALW and ESPript) of PsaL among *Cyanophora* and other photosynthetic organisms.**

The amino acid residues of *Cyanophora* PsaL that cause steric hindrance with the cyanobacterial trimer are highlighted in cyan. The purple boxes show regions at the N-terminus and C-terminus that are lacking in *Cyanophora*. Completely conserved residues are highlighted in red. The species shown are *Cyanophora paradoxa*, *Thermosynechococcus elongatus* BP-1, *Synechocystis* sp. PCC 6803, *Anabaena* sp. PCC 7120, *Chaetoceros gracilis*, *Chlamydomonas reinhardtii*, *Cyanidioschyzon merolae*, *Pisum sativum*, and *Zea mays*.

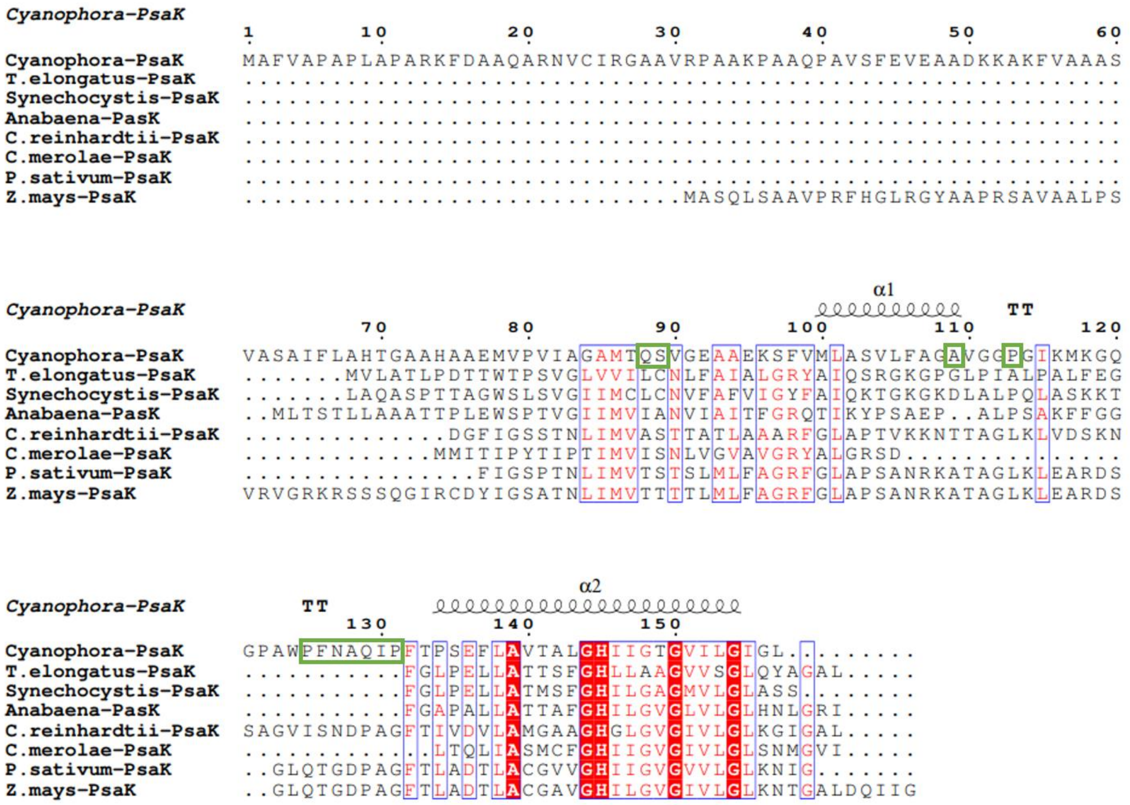

**Supplementary Figure 16. Multiple sequence alignment (CLUSTALW and ESPript) of PsaK among *Cyanophora* and other photosynthetic organisms.**

The amino acid residues in the contact regions at the center of the PSI tetramer are highlighted in green. Completely conserved residues are highlighted in red. The species shown are *Cyanophora paradoxa*, *Thermosynechococcus elongatus* BP-1, *Synechocystis* sp. PCC 6803, *Anabaena* sp. PCC 7120, *Chlamydomonas reinhardtii*, *Cyanidioschyzon merolae*, *Pisum sativum*, and *Zea mays*.

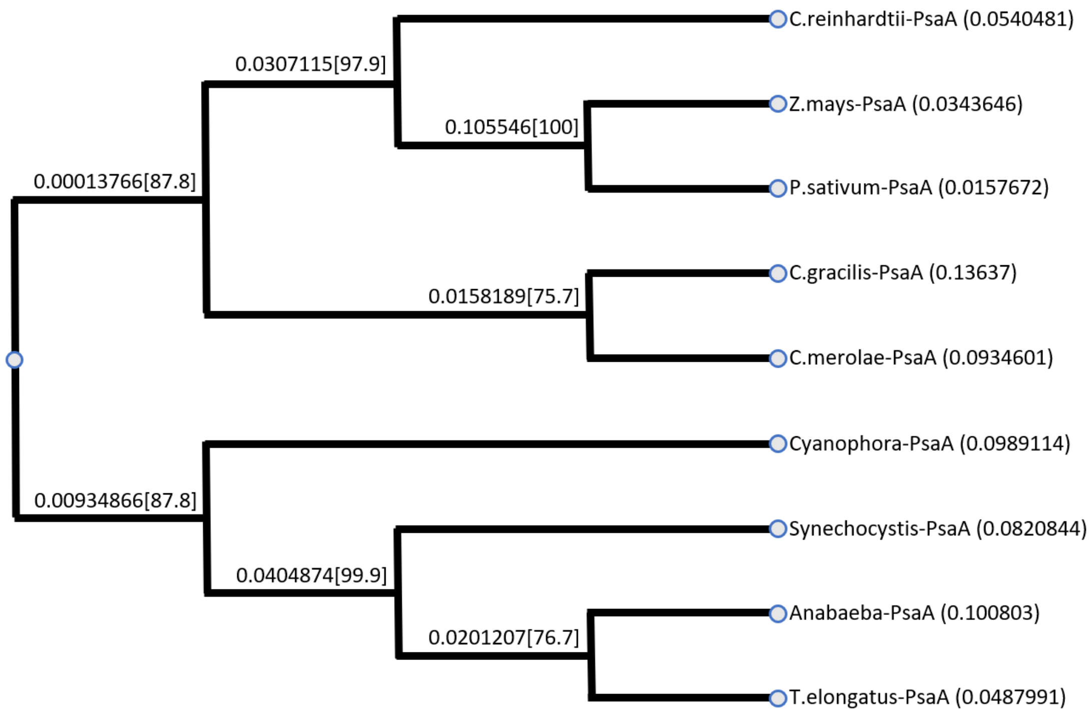

**Supplementary Figure 17. Phylogenies analysis of PsaA among *Cyanophora* and other photosynthetic organisms.**

The species shown are *Cyanophora paradoxa*, *Thermosynechococcus elongatus* BP-1, *Synechocystis* sp. PCC 6803, *Anabaena* sp. PCC 7120, *Chaetoceros gracilis*, *Chlamydomonas reinhardtii*, *Cyanidioschyzon merolae*, *Pisum sativum*, and *Zea mays*.

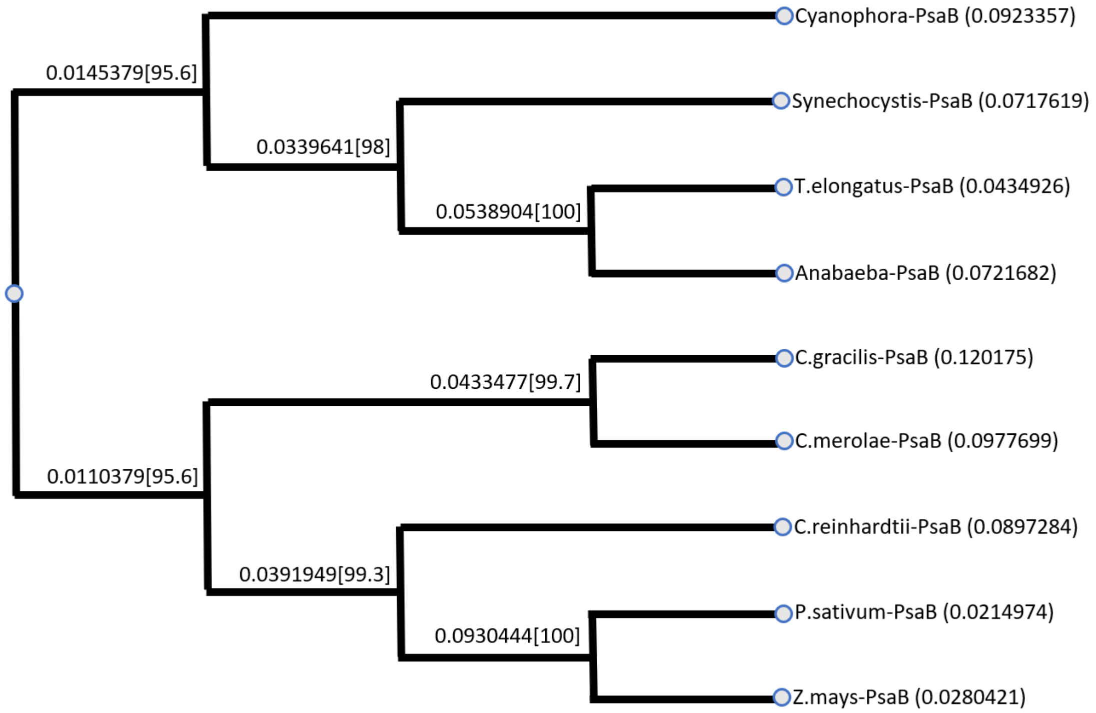

**Supplementary Figure 18. Phylogenies analysis of PsaB among *Cyanophora* and other photosynthetic organisms.**

The species shown are *Cyanophora paradoxa*, *Thermosynechococcus elongatus* BP-1, *Synechocystis* sp. PCC 6803, *Anabaena* sp. PCC 7120, *Chaetoceros gracilis*, *Chlamydomonas reinhardtii*, *Cyanidioschyzon merolae*, *Pisum sativum*, and *Zea mays*.

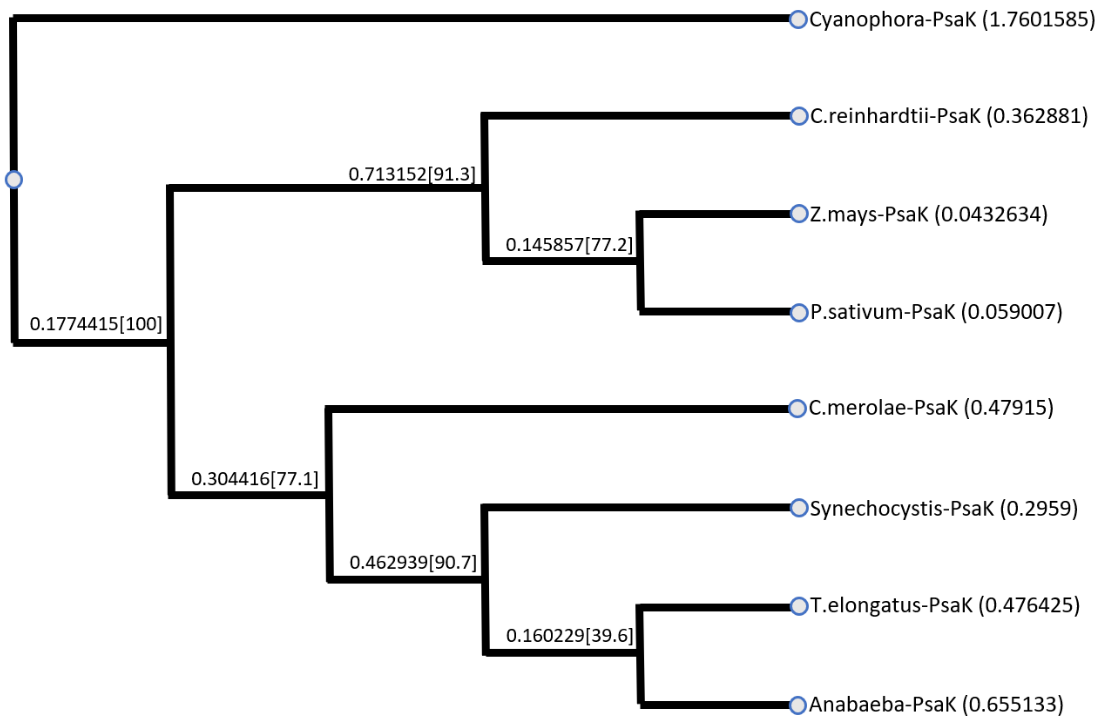

**Supplementary Figure 19. Phylogenies analysis of PsaK among *Cyanophora* and other photosynthetic organisms.**

The species shown are *Cyanophora paradoxa*, *Thermosynechococcus elongatus* BP-1, *Synechocystis* sp. PCC 6803, *Anabaena* sp. PCC 7120, *Chlamydomonas reinhardtii*, *Cyanidioschyzon merolae*, *Pisum sativum*, and *Zea mays*.

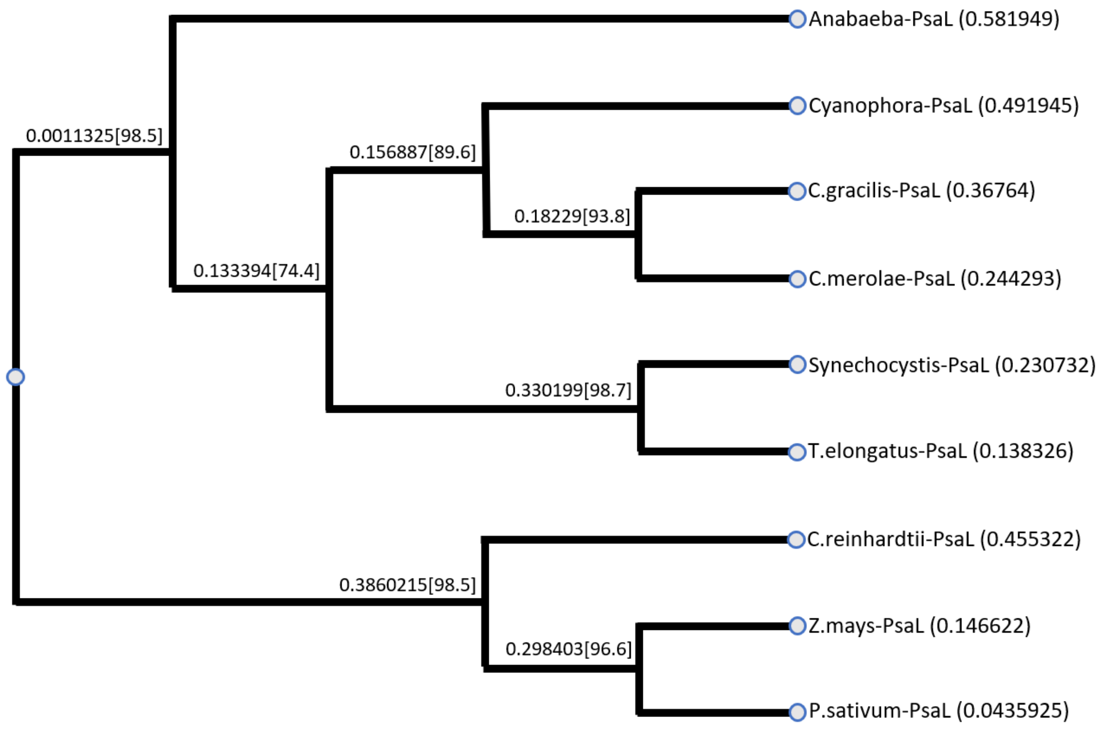

**Supplementary Figure 20. Phylogenies analysis of PsaL among *Cyanophora* and other photosynthetic organisms.**

The species shown are *Cyanophora paradoxa*, *Thermosynechococcus elongatus* BP-1, *Synechocystis* sp. PCC 6803, *Anabaena* sp. PCC 7120, *Chaetoceros gracilis*, *Chlamydomonas reinhardtii*, *Cyanidioschyzon merolae*, *Pisum sativum*, and *Zea mays*.

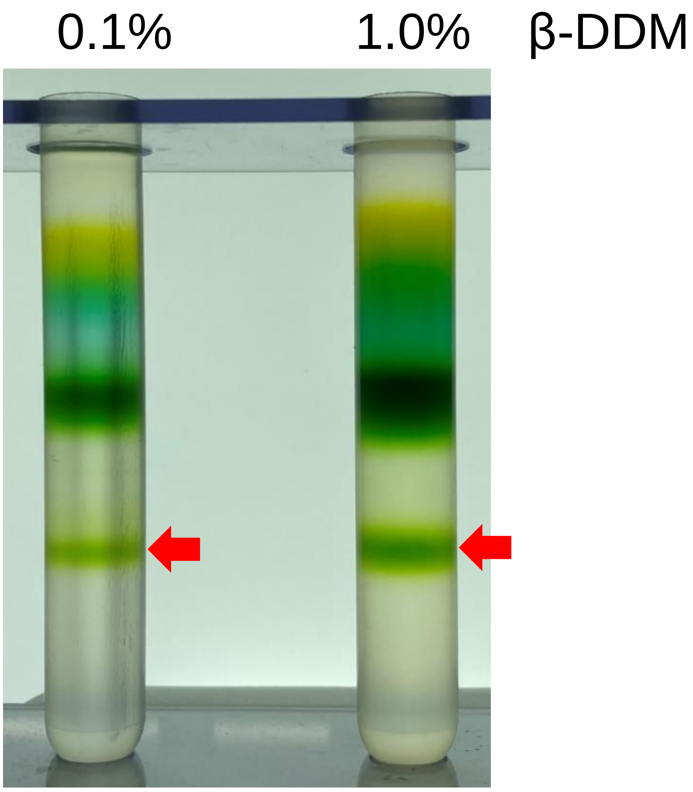

**Supplementary Figure 21. Trehalose gradient centrifugation after solubilizing the thylakoids with 0.1% or 1.0% *β*-DDM.**

Red arrows indicate the PSI tetramer. Thylakoid membranes were solubilized with either 0.1% or 1% *β*-DDM at a Chl concentration of 0.25 mg $mL^{-1}$ for 30 min on ice in the dark with gentle stirring. After centrifugation at 20,000 × g for 10 min at 4°C, the resultant supernatant were concentrated using a 100 kDa cut-off filter (Amicon Ultra; Millipore) at 4,000 × g, and then the concentrated samples were loaded onto a linear trehalose gradient of 10–40% (w/v) in a medium containing 20 mM Mes-NaOH (pH 6.5), 0.2 M NaCl, and 0.1% *β*-DDM. The PSI-tetramer fraction was obtained after centrifugation at 154,000 × g for 18 h at 4°C.

**Supplementary Table 1. Statistics of data collection, processing, and refinement.**

| Complex | Wild-type PSI monomer1 | Wild-type PSI monomer2 | Wild-type PSI tetramer | GraFix PSI tetramer |
|---|---|---|---|---|
| PDB ID | 7DR0 | 7DR1 | - | 7DR2 |
| EMDB ID | EMD-30820 | EMD-30821 | EMD-30822 | EMD-30823 |
| Data collection and processing | | | | |
| Microscope | | Talos Arctica | | |
| Detector | | Falcon 3EC direct electron detector | | |
| Magnification | | 92000 | | |
| Voltage (kV) | | 200 | | |
| Defocus range (μm) | | -2.0 to -4.0 | | -1.5 to -3.5 |
| Pixel size (Å) | | 1.093 | | |
| Total electron dose ($e^-/Å^2$) | | 50 | | |
| Exposure time (s) | | 3.7 and 4.0 | | 3.2 |
| Number of frames per image | | 37 and 40 | | 32 |
| Number of micrographs | | 4,515 | | 3,205 |
| Initial particle images (no.) | | 1,603,082 | | 724,316 |
| Final particle images (no.) | 70,920 | 110,380 | 145,567 | 40,679 |
| Map resolution (Å) | 3.3 | 3.2 | 4.0 | 3.8 |
| Applied b-factor ($Å^2$) | -107 | -117 | -216 | -168 |
| Applied symmetry | C1 | C1 | C2 | C2 |
| Refinement | | | | |
| Initial model used | Homology model | Homology model | - | Homology model |
| Model resolution (Å) | 3.3 | 3.2 | - | 3.8 |
| FSC threshold | 0.5 | 0.5 | - | 0.5 |
| No. of atoms | | | | |
| Protein | 16,975 | 16,572 | - | 67,224 |
| Ligand | 5,771 | 5,606 | - | 22,732 |
| B factors ($Å^2$) | | | | |
| Protein | 44.0 | 35.0 | - | 69.5 |
| Ligand | 34.3 | 26.2 | - | 66.3 |
| R.m.s deviations | | | | |
| Bond lengths (Å) | 0.009 | 0.007 | - | 0.009 |
| Bond angles (°) | 1.50 | 1.53 | - | 1.67 |
| Validation | | | | |
| MolProbity score | 1.91 | 1.84 | - | 2.12 |
| Clashscore | 9.16 | 8.09 | - | 11.5 |
| Poor rotamers (%) | 0.62 | 0.75 | - | 1.27 |
| EMRinger score | 3.4 | 3.6 | - | 1.58 |
| Ramachandran plot | | | | |
| Favored (%) | 93.62 | 93.98 | - | 92.58 |
| Allowed (%) | 6.15 | 5.74 | - | 7.16 |
| Disallowed (%) | 0.23 | 0.29 | - | 0.26 |

**Supplementary Table 2. Cofactors in each monomer unit of the *Cyanophora* PSI tetramer and comparison with a monomer of *T. elongatus* PSI trimer.**

| Protein | | Chlorophyll | Carotenoid | Lipid | Others |
|---|---|---|---|---|---|
| PsaA | monomer1 | 46 Chl *a* | 6 BCR | 2 LHG | 1 [4Fe-4S] cluster, 1 phylloquinone |
| | monomer2 | 45 Chl *a* | 6 BCR | 2 LHG | 1 [4Fe-4S] cluster, 1 phylloquinone |
| | *T. elongatus* PSI† | 46 Chl *a* | 6 BCR | 2 LHG | 1 [4Fe-4S] cluster, 1 phylloquinone |
| PsaB | monomer1 | 32 Chl *a* | 4 BCR | 1 LMG | 1 phylloquinone |
| | monomer2 | 31 Chl *a* | 4BCR | 1 LMG | 1 phylloquinone |
| | *T. elongatus* PSI† | 41 Chl *a* | 7 BCR | 1 LMG<br>1 LHG | 1 phylloquinone |
| PsaC | monomer1 | | | | 2 [4Fe-4S] cluster |
| | monomer2 | | | | 2 [4Fe-4S] cluster |
| | *T. elongatus* PSI† | | | | 2 [4Fe-4S] cluster |
| PsaD | monomer1 | | | | |
| | monomer2 | | | | |
| | *T. elongatus* PSI† | | | | |
| PsaE | monomer1 | | | | |
| | monomer2 | | | | |
| | *T. elongatus* PSI† | | | | |
| PsaF | monomer1 | 1 Chl *a* | 1 BCR | | |
| | monomer2 | 1 Chl *a* | 1 BCR | | |
| | *T. elongatus* PSI† | 1 Chl *a* | 1 BCR | | |
| PsaI | monomer1 | | 2 BCR | | |
| | monomer2 | | 2 BCR | | |
| | *T. elongatus* PSI† | | 2 BCR | | |
| PsaJ | monomer1 | 1 Chl *a* | 3 BCR | | |
| | monomer2 | 1 Chl *a* | 3 BCR | | |
| | *T. elongatus* PSI† | 2 Chl *a* | 3 BCR | | |
| PsaK⁋ | monomer1 | 1 Chl *a* | | | |
| | monomer2 | | | | |
| | *T. elongatus* PSI† | 1 Chl *a* | | | |
| PsaL | monomer1 | 3 Chl *a* | 2 BCR | | |
| | monomer2 | 3 Chl *a* | 2 BCR | | |
| | *T. elongatus* PSI† | 3 Chl *a* | 2 BCR | | |
| PsaM | monomer1 | | 1 BCR | | |
| | monomer2 | | 1 BCR | | |
| | *T. elongatus* PSI† | 1 Chl *a* | 1 BCR | | |
| PsbX⁋ | monomer1 | | | | |
| | monomer2 | | | | |
| | *T. elongatus* PSI† | 1 Chl *a* | | | |
| Total | monomer1 | 84 | 19 | 3 | 5 |
| | monomer2 | 81 | 19 | 3 | 5 |
| | *T. elongatus* PSI† | 96 | 22 | 4 | 5 |

BCR, $\beta$-carotene; LMG, distearoylmonogalactosyl diglyceride; LHG, dipalmitoylphosphatidyl glycerol.

†PDB: 1JB0.

⁋PsaX in monomer1 and PsaK and PsaX in monomer2 are not found in the *Cyanophora* PSI.

**Supplementary Table 3. Chls lost in the *Cyanophora* PSI tetramer and their ligands.**

| | *T. elongatus* PSI[†] Chls[¶] | *T. elongatus* PSI[†] Ligands | *Cyanophora*-monomer1 Ligands | *Cyanophora*-monomer2 Ligands |
|---|---|---|---|---|
| PsaA | 1402 | - | (Chl845)** | - |
| PsaB | 1209 | H176 | H177 | H177 |
| | 1216 | $H_2O$ | - | - |
| | 1217 | H288 | H289 | H289 |
| | 1218 | H298 | H299* | H299* |
| | 1219 | $H_2O$ | - | - |
| | 1220 | H322 | H319* | H319* |
| | 1221 | Y329 | (V326) | (V326) |
| | 1227 | H417 | H414 | H414 |
| | 1232 | $H_2O$ | (Chl835)** | - |
| | 1233 | $H_2O$ | - | - |
| PsaJ | 1303 | H39 | (Y38) | (Y38) |
| PsaK | 1401 | - | (Chl201)** | - |
| PsaM | 1601 | $H_2O$ | - | - |
| PsaX | 1701 | N23 | - | - |

[†]PDB: 1JB0.

[¶]These Chls listed cannot be seen in the *Cyanophora* PSI structure compared with the *T. elongtus* PSI structure.

*Disorder.

**Existing in monomer1.
